# Climate-driven variation in biotic interactions provides a narrow and variable window of opportunity for an insect herbivore at its ecological margin

**DOI:** 10.1101/2021.10.27.466153

**Authors:** James E. Stewart, Ilya M.D. Maclean, Gara Trujillo, Jon Bridle, Robert J. Wilson

**Affiliations:** College of Life and Environmental Sciences, University of Exeter, United Kingdom; Environment & Sustainability Institute, University of Exeter, Penryn Campus, United Kingdom; International Institute for Industrial Environmental Economics (IIIEE), Lund University P.O. Box 196, 22100 Lund, Sweden; School of Biological Sciences, University of Bristol, United Kingdom; Department of Genetics, Evolution, and Environment, University College London, London; Departmento de Biogeografía y Cambio Global, Museo Nacional de Ciencias Naturales, Madrid E28006, Spain

**Keywords:** brown argus, Lepidoptera, host shift, specialisation, asynchrony

## Abstract

Climate-driven geographic range shifts have been associated with transitions between dietary specialism and generalism at range margins. The mechanisms underpinning these often transient niche breadth modifications are poorly known, but utilisation of novel resources likely depends on phenological synchrony between the consumer and resource. We use a climate-driven range and host shift by the butterfly *Aricia agestis* to test how climate-driven changes in host phenology and condition affect phenological synchrony, and consider implications for host use.

Our data suggest that the perennial plant which was the primary host before range expansion is a more reliable resource than the annual Geraniaceae upon which the butterfly has become specialised in newly colonised parts of its range. In particular, climate-driven phenological variation in the novel host *Geranium dissectum* generates a narrow and variable ‘window of opportunity’ for larval productivity in summer. Therefore, although climatic change may allow species to shift hosts and colonise novel environments, specialisation on phenologically-limited hosts may not persist at ecological margins as climate change continues. We highlight the potential role for phenological (a)synchrony in determining lability of consumer-resource associations at range margins, and the importance of considering causes of synchrony in biotic interactions when predicting range shifts.

## Introduction

Climate change is causing widespread shifts in species’ geographic range limits [1–3]. The extent of such shifts depends on species’ life histories, potential for plastic responses and the quality of available habitat at the expanding margin [4–6]. Habitat availability may itself be determined by the process of range expansion: range shifts have recently been identified as a cause, rather than consequence, of increased dietary generalism at poleward range margins in herbivorous insects [7,8]. However, dietary generalism and incorporation of novel host plants can be transient in such systems, and the mechanisms underlying gain or loss of hosts from insect diets is poorly known [7–9]. Here, we highlight phenological (a)synchrony between insects and unpredictable host resources as a potential mechanism for lability in insect-host associations at range margins.

The phenology of many consumers is closely synchronised with the development and availability of their resources, and their interactions occur within an often narrow ‘window of opportunity’ for the consumer defined by the phenology of both partners [10–14]. For herbivorous insects, the length of the phenological window of opportunity will be partly determined by the specificity of its interaction with the host, the host’s growth form, and the broader environmental context. For example, the window of opportunity for herbivory is typically longer on perennials than on short-lived annual plants, the availability of which may be defined by environmental drivers of their germination and senescence [15,16]. The window is also typically longer for polyphagous than obligately monophagous species (which can exploit fewer distinct phenological windows), and for populations inhabiting topographically variable landscapes in which heterogeneous microclimates provide diverse phenological windows [16,17]. Differences in synchrony among nearby microclimates may also cause local variation in host condition, quality and profitability for the herbivore, thereby influencing local patterns of host selection and opportunities for dietary change [7,8,18–20].

Robust evidence is therefore required on the drivers and vulnerability of host-herbivore phenological synchrony (which may scale up to emergent patterns of herbivore range and host shifts [10,19,21]), including the role of microclimate in consumer persistence by potentially buffering asynchrony in biotic interactions [18,20]. Here we address this knowledge gap, using as an exemplar the brown argus butterfly *(Aricia agestis;* Lepidoptera: Lycaenidae) at its range margin in the UK. In doing so, we highlight how phenological (a)synchrony could provide an underlying mechanism for apparently high rates of host shifting near range margins [7,22]. Brown argus butterflies have two generations per year; larval offspring of the second generation emerge from mid-August and feed on leaves of the host plant before overwintering [23]. The brown argus’ UK range was historically largely restricted to calcareous grassland where its host, the perennial *Helianthemum nummularium* (Malvales: Cistaceae; hereafter *Helianthemum),* grows. However, since the 1990s, the brown argus has undergone a climate-driven range expansion associated with rapid evolution of biotic interactions, including specialisation on annual Geraniaceae species (*Erodium cicutarium*, *Geranium dissectum* and *G. molle*; Geraniales: Geraniaceae) mainly in regions beyond the former range limit [21,24–27].

In this study, we test expectations of (a) greater temporal variation in the condition of the annual versus perennial host plants, and (b) more opportunity for asynchrony between the consumer and its annual hosts than its perennial host, as a consequence of (a). We expect asynchrony with the less-predictable hosts to be more pronounced under warm, dry summer conditions, and we test for such effects on the condition and phenology of the annual Geraniaceae hosts that have enabled the range expansion. We conduct these tests (c) across sites and years, and (d) across microclimates within a site. We then consider the implications of asynchrony for host use, shifting host associations at range margins, and the range dynamics of host-limited herbivores.

## Methods

### Study system

Brown argus butterflies prefer to lay their eggs on Geraniaceae, on which larvae grow 10% larger and faster, than on *Helianthemum* (the ancestral host at the range margin in Britain), and prefer to lay on better condition leaves regardless of host species [19,21,24–26,28,29]. However, the annual, more ephemeral growth form of the Geraniaceae host plants may make them less reliable as a food source than the evergreen perennial *Helianthemum,* especially under more variable climatic conditions at the range margin [16]. To investigate this, we surveyed ten sites fortnightly–monthly between July 2016 and October 2017, to monitor phenology and condition of three host plant species. *Helianthemum* was the dominant host at five calcareous grassland sites, while Geraniaceae (*G. dissectum* and *E. cicutarium*; hereafter *Geranium* and *Erodium*) were dominant at five grassland/ dune sites (Figure 1). The number of sites and quadrats was chosen to maximise spatial coverage and replication within logistical constraints. See SI-1.1 and SI-1.2 for survey dates and site profiles.

**Figure 1.**
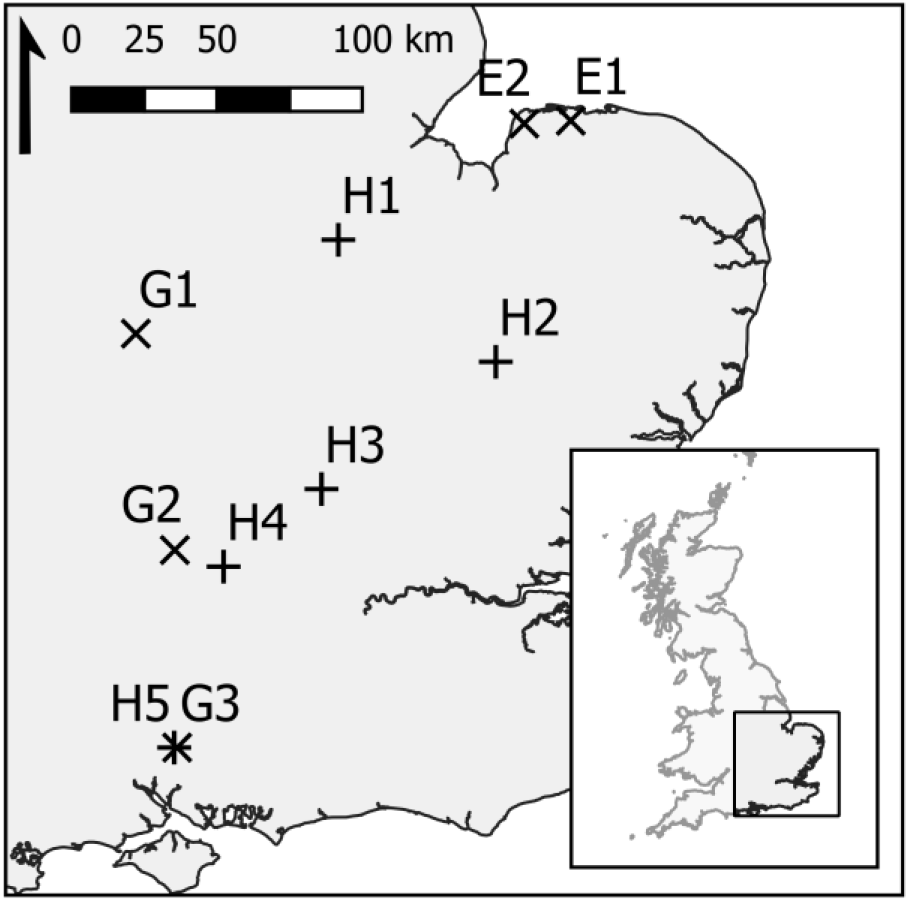
Map of study sites in England. + denotes *Helianthemum* sites (H1–H5, Table S1), × denotes Geraniaceae sites (*Geranium*, G1–G3; *Erodium,* E1–E2; Table S1).

### Quadrat surveys and analyses

Using an average of 33.7 (0.25 m^2^) quadrats per survey (SI-1.3 for quadrat placement), we recorded percentage cover (SI-1.3.2), phenophase (phenological stage) and condition of each host species. Host phenophase and condition underpin brown argus egg-laying site choice in the field [19,30]. Phenophase was estimated on a four-point scale (in leaf L, bud B, flower F, or had set seed S). Condition was visually assessed on a scale of 0–3 (poor–high quality for egg-laying, following [19,26]; see SI-1.3.3 for details and justification). We also measured mean sward height and percentage cover of bare ground (SI-1.3) which can alter local microclimates [31,32].

*Quadrat-level* phenophase and condition were estimated based on the average for each host species in each quadrat. This approach can mask fine-scale changes in phenophase and host condition, particularly for the annual hosts, so we also recorded *plant-level* phenophase and condition of the smallest, earliest phenophase Geraniaceae plant in each quadrat. This approach allows inference as to whether new germination has occurred, and of the age/condition of the plant material likely to be available to overwintering larvae. By September, Geraniaceae plants in later phenophases are typically senescent, in poor condition and expected to die before the autumn, making them a poor resource for larvae [30].

### Quantifying variation in host plant condition and phenophase

To test expectation (a), for greater temporal variation in the condition of the annual than perennial hosts, we used a Kruskal-Wallis (KW) test for each sampling period to compare quadrat-level condition between host species. Bonferroni-corrected Dunn’s tests were then used to identify which hosts differed significantly from one another in condition score within each sampling period. We then conducted interannual comparisons of Geraniaceae host condition (Mann-Whitney U tests, MWU) and phenophase (*X*^2^ tests): these compared 2016 data with 2017 data, separately for each month between July to October. These months are the most relevant for host choice and larval feeding by second generation brown argus and their offspring.

### Assessing plant-herbivore (a)synchrony

To contextualise host condition with reference to herbivore phenology, and address question (b), we overlaid plots of site-specific host condition indices (calculated using survey data, SI-1.5.1) from all July–October surveys with emergence phenology curves of adult second generation brown argus and their larval offspring. Adult phenology was described as site-specific Gaussian curves for 2016 and 2017, based on output of phenomenological models following [30,33] (below and SI-1.6). The larval phenology curves track the adult curves, with an estimated 11-day lag to account for mating and egg-laying (four days post-emergence) and larval emergence (one week) [24]. Therefore, the larval emergence curves are presented as indicators rather than precise evaluations of appearance or abundance at each site.

The overlap between plotted condition indices and brown argus phenology curves was used to generate an area under the curve (AUC) metric of site- and year-specific synchrony between brown argus and the host plants (SI-1.7). These AUC metrics were then modelled in a beta regression (logit link; SI-1.7) to test for effects of site latitude, host plant, year and a host-year interaction on synchrony.

The phenomenological models used to generate phenology estimates account for variation in phenology between sites and years based on differences in latitude and temperature (SI-1.6). In summary, brown argus second brood phenology in Britain varies with latitude (earlier further north) and between-brood temperature (earlier under warmer conditions between the first and second brood), and is related to the (latitude-dependent) phenology of the first brood [30,33]. See SI-1.6 for more information.

### Testing climatic predictors of Geraniaceae recruitment and condition

To address our third question (c), using data on the youngest host within each quadrat, we tested for climatic drivers of recruitment and condition of each Geraniaceae species in early September, when most larval offspring were expected to have emerged to feed. We defined recruitment as the presence of at least one young, leaf-stage host plant in the focal quadrat. Plants in condition categories 1 and 2 were rarely observed during September surveys. We therefore reclassified condition 0/1 plants as poor condition (0) and condition 2/3 plants as good condition (1).

To test predictors of recruitment and condition, we used logistic regression with the following putative predictors: year, site, northing, easting, vegetation height, bare ground cover, and (linear and quadratic terms for) local weather estimates based on the UK Meteorological Office’s 5 km gridded weather data [34]. Using daily weather data [34], we calculated the minimum, mean and maximum temperature and mean rainfall for three periods in each year (justified in SI-1.7): July, July–September and August–September (only including weather data up to the day of quadrat sampling at each site in early September). We also used daily weather data to calculate the Gaussen Aridity Index (GAI; precipitation/(2 × temperature)) for each period (e.g. [35]). Higher GAI values indicate cooler, wetter conditions. Day of year was tested both as a putative fixed effect predictor and as an offset term to account for day of sampling.

All continuous predictors were standardised and site (*n* ≤ 5 per host plant) was included as a fixed effect (following [36]). We constructed candidate models by considering all plausible parameter combinations (including temperature-rainfall interactions), estimated parameters using maximum likelihood, and used AIC-based model selection to establish the most parsimonious model(s) (see SI-1.9–1.10 for details of model selection, validation and diagnostics).

### Testing microclimate effects on Geraniaceae phenology and condition

In September 2017, we calculated condition and phenophase indices (SI-1.5.2) for each of 31 quadrats at site G1, using the phenophase (L, B, F and S) and condition (0–3) of all *Geranium* plants in each quadrat. The indices range between 0 (quadrats contain only plants at condition 0/seed set stage) and 1 (plants at condition 3/leaf stage). To test expectation (d), we modelled the indices (logistic regression) as a function of putative quadrat-specific predictors: microclimate (mean, maximum and minimum temperature and soil moisture; SI-1.11), percentage cover of bare ground and mean sward height. We also considered plausible temperature-moisture interactions.

## Results

### Variation in host plant condition and phenophase

There was a substantial decline in quadrat-level condition of the annual host *Geranium* over summer 2016 that was not observed in the perennial *Helianthemum* or to the same extent in *Erodium* (Figure 2a; KW tests, Table S4). However, by early November 2016, senesced *Geranium* had mostly been replaced by recently-germinated, better condition recruits, and the quadrat-level condition was at least as high as that of *Helianthemum* (Figure 2a-c; KW tests, Table S4).

**Figure 2.**
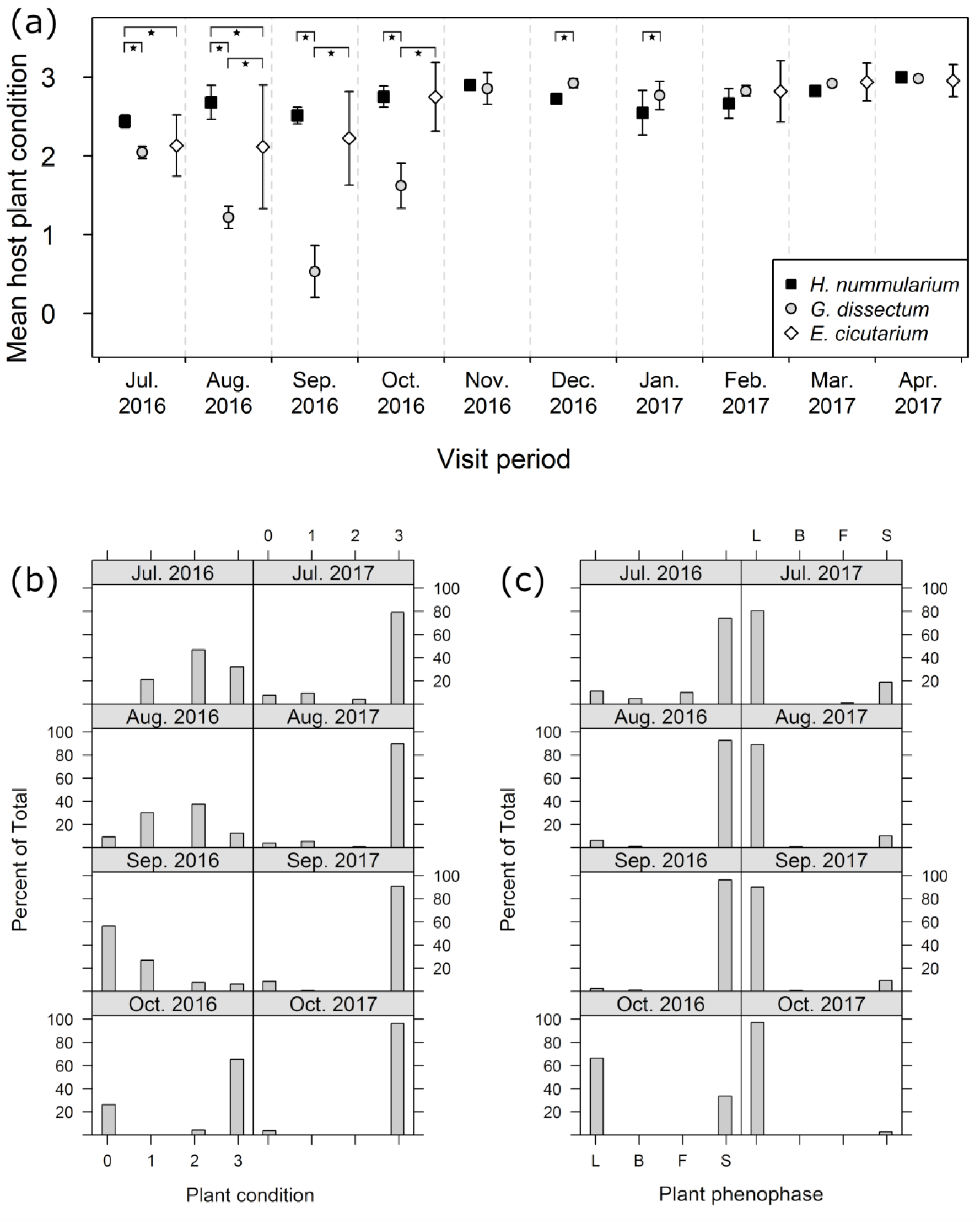
(a) Mean (± SD) of site-averaged quadrat-level host plant condition categorisations (0–3). Starred brackets indicate significant differences in condition between hosts (Table S4). Condition (0–3; b) and phenophase (c) of the youngest *G. dissectum* plant in each quadrat across all sites visited during late July–early October in both 2016 and 2017. Phenophase L: leaf; B: in bud; F: in flower; S: set seed. Equivalents of (b) and (c) for *Erodium* are available as Figure S3.

Overall, in summer 2016 both Geraniaceae species showed substantial evidence of leaf senescence, including wilting and abscission, and little germination or seedling establishment until early October *(pers. obs.) (Geranium,* Figure 2b-c; *Erodium,* Figure S3). By contrast, in 2017, there were many more seedlings and good condition plants throughout the summer months and into the main sampling period in September *(pers. obs.)* (Geranium, Figure 2b-c; *Erodium*, Figure S3).

The condition of the youngest *Geranium* in each quadrat was significantly higher in all 2017 survey periods compared to 2016 (Figure 2b; MWU tests, Table S5). *Geranium* phenophases differed significantly between years *(χ^2^* tests, Table S5): in 2016, the youngest plant in each *Geranium* quadrat was typically an older plant in the seed set stage (Figure 2c) and new plants in the leaf stage did not dominate until October; however, this younger form was dominant throughout summer 2017 (Figure 2c).

*Erodium* showed similar patterns, though condition was approximately equivalent between years for the August and September surveys (Figure S3; Mann-Whitney U tests, Table S5). *Erodium* quadrats were also dominated by young, leaf-stage plants in 2017: significantly more so than in 2016 during July and August (Figure S3; *χ*^2^ tests, Table S5). Similar patterns of Geraniaceae phenophase and condition were observed at quadrat-level as these plant-level assessments (Figures S4 and S5).

### Plant-herbivore (a)synchrony

We assessed potential for asynchrony between the butterfly and its hosts (question (b)) by overlaying plots of site-specific host condition indices with curves representing brown argus phenology (Figure 3), and performing beta regressions on derived synchrony estimates. The beta regressions of AUC synchrony estimates demonstrate that synchrony was lowest for brown argus on *Geranium* in 2016 (low AUC overlap: AUC range 0.51–0.73), but very high for other hosts and for *Geranium* in 2017 (high AUC overlap: AUC range 0.92– 1.00) (Figure 3, Tables 1 and S6). Synchrony was lower for larvae than adults, especially on *Geranium* in 2016 (Table S6). There was no detectable effect of site latitude on adult or larval AUC overlap (SI-1.12.4).

**Figure 3.**
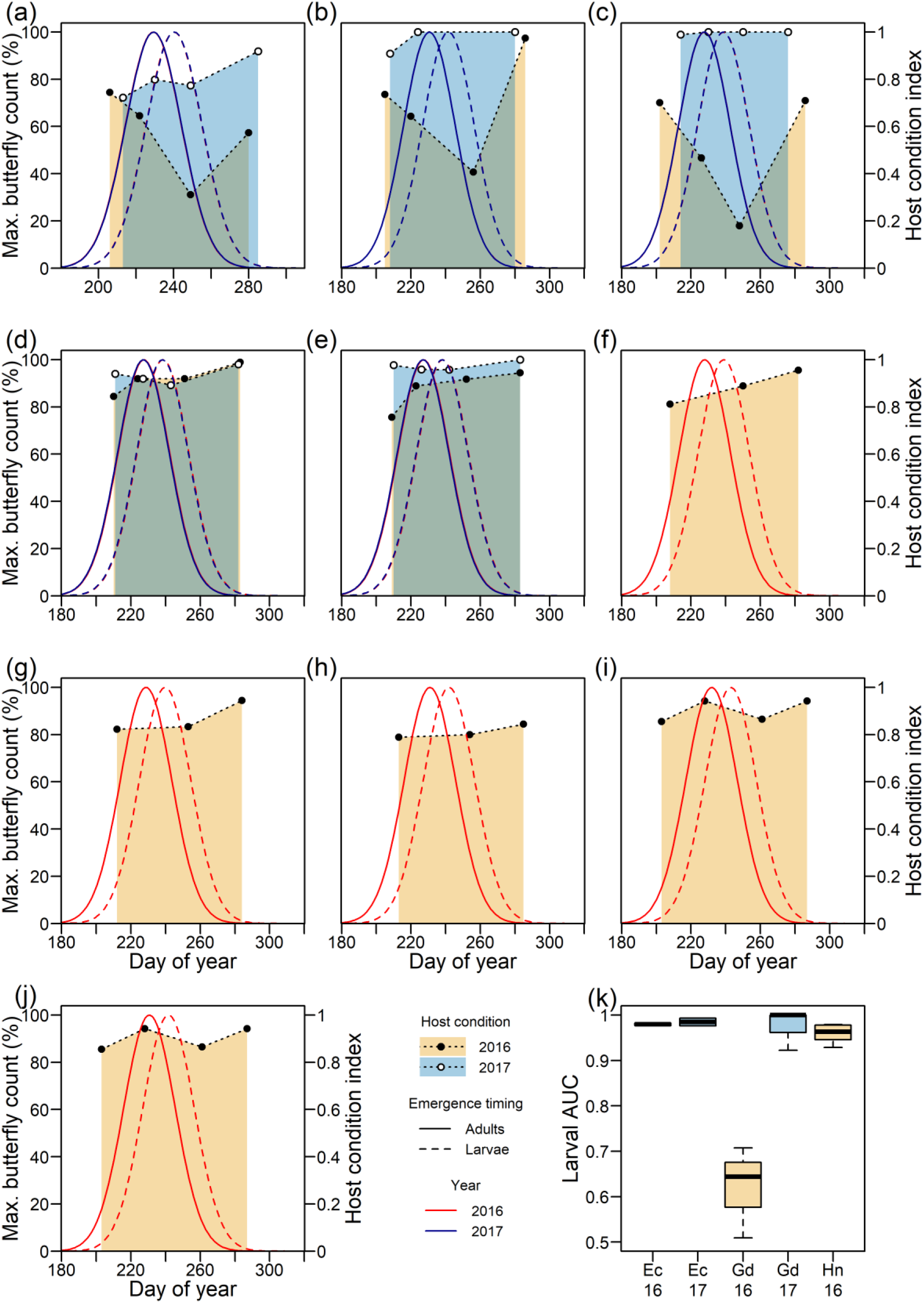
Plant condition index in 2016 and 2017 for *Geranium* at sites G1–G3 (a–c), *Erodium* at sites E1–E2 (d–e), and *Helianthemum* at sites H1–H5 (f–j), indicating timing of host condition changes relative to the year- and site-specific emergence of second generation brown argus adults and their larvae. Butterfly phenology curves typically overlap at each site. (k) summarises larval AUC synchrony metrics for each host-year combination, summarised from site-specific metrics each calculated as the full area of the phenology curve minus that which lies above the corresponding host condition line. The equivalent plot for adults is shown in SI-1.12.4.

**Table 1.** Parameter estimates (with errors) for Bayesian beta regression of adult and larval AUC synchrony metrics, also showing the lower (L95) and upper (U95) credible intervals for each estimate. Predictors include year, host and host-year interaction.

|  | Adults |  |  | Larvae |  |  |
| --- | --- | --- | --- | --- | --- | --- |
|  | Estimate | L95 | U95 | Estimate | L95 | U95 |
| **Intercept** | 2.89 (0.44) | 2.03 | 3.77 | 2.8 (0.4) | 1.99 | 3.55 |
| **Year<sub>2017</sub>** | 1.61 (0.65) | 0.38 | 2.93 | 1.69 (0.65) | 0.46 | 3.01 |
| **Host<sub>Geranium</sub>** | -2.01 (0.48) | -2.94 | -1.07 | -2.23 (0.45) | -3.07 | -1.32 |
| **Host<sub>Helianthemum</sub>** | 0.02 (0.48) | -0.94 | 0.97 | 0.01 (0.37) | -0.7 | 0.73 |
| **Year<sub>2017</sub>:Host<sub>Geranium</sub>** | 2.87 (0.73) | 1.43 | 4.29 | 3.06 (0.74) | 1.58 | 4.5 |

Estimates for *Erodium* and Year<sub>2016</sub> are not shown here as they are the base level for each predictor and are therefore included in the intercept term.

In 2016, *Geranium* condition declined over the peak of adult brown argus emergence, and was lowest during the period in which most larvae would be beginning to feed (Figure 3a–c). By contrast, at their respective sites, good-condition *Erodium* and *Helianthemum* hosts were available throughout egg-laying and early larval feeding periods of the brown argus butterfly, with little variation between two climatically different years (2016 and 2017), or among sites at each sampling period (Figure 3d–j). Among-site variation in condition is more pronounced for *Geranium,* but this does not mask the temporal variation within and between years (Figure 3; Table S4). Among-site variation likely results from local variation in temperature and water relations linked to factors including weather, topography and geology.

### Climatic predictors of Geraniaceae condition and recruitment

We integrated climate data with in-field host plant surveys to investigate potential drivers of condition and phenology across the *Geranium* sites, addressing question (c). The probability of the youngest *Geranium* plants being in good condition increased with summer rainfall (*MC_final_*; Figure 4a, Table 2a). There was limited evidence that moister, cooler conditions in areas with shorter vegetation were associated with better condition (Table S8). Forcing models of *Geranium* condition to include an effect of year resulted in higher AIC values, and inflated parameter estimates and standard errors by several orders of magnitude, so these are reported only in the Supplementary Information for context (SI-1.12.5, Table S8), and there are no effects of year in the final model set. The evidence suggests that probability of new *Geranium* recruitment was higher in 2017 (*MR_best_*; SI-1.12.5, Table S9), and lower following higher mean daily temperatures during August–September (*MR_final_*; Figure 4b, Table 2b). There were no detectable effects of site northing or easting, or day of sampling on condition or recruitment.

**Figure 4.**
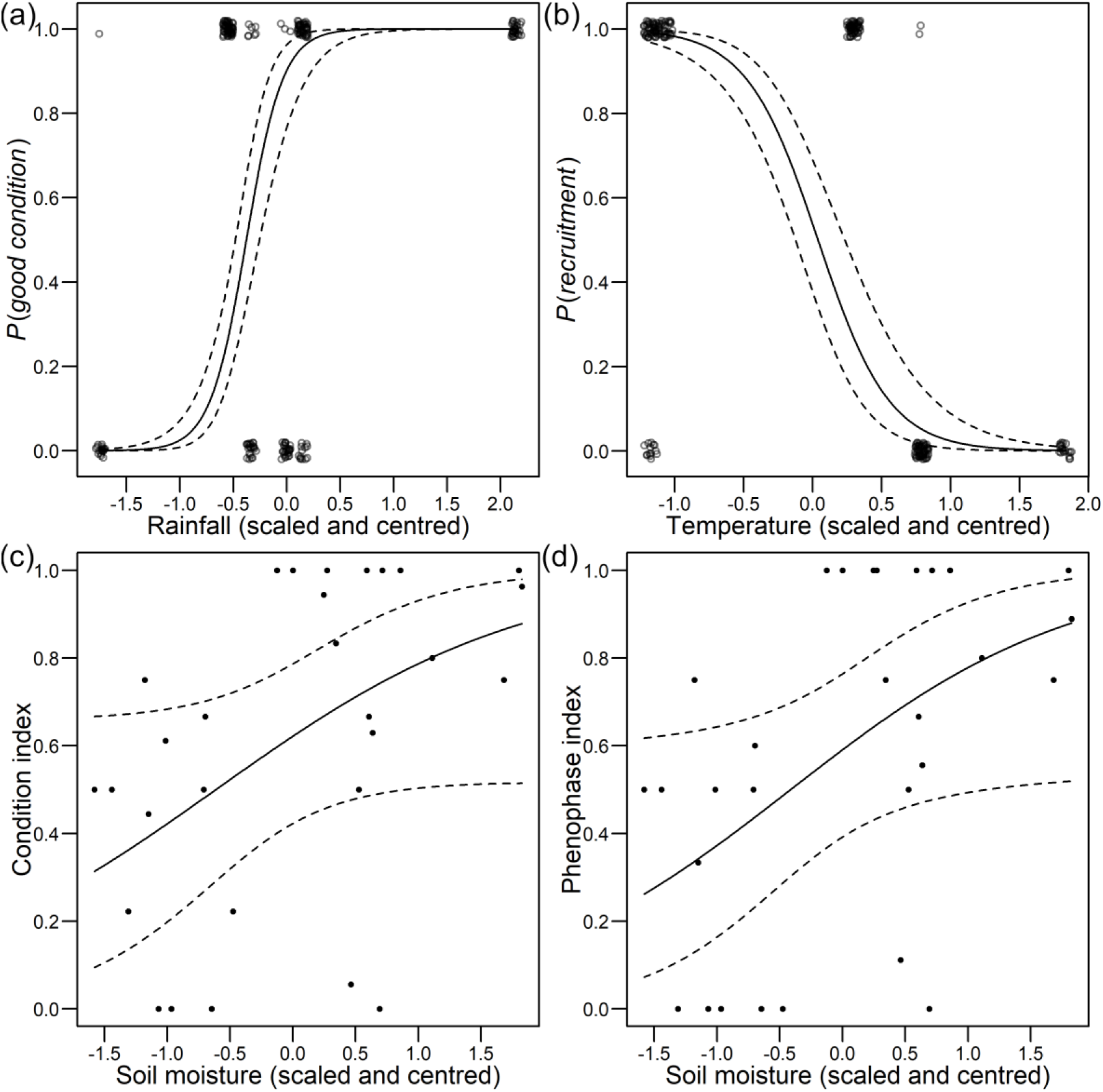
(a,b) Predicted probabilities with 95% confidence intervals (dashed lines) of (a) a state of good condition in the youngest *Geranium* in each quadrat, and (b) recruitment of *Geranium* in each quadrat. The probabilities (a) increase as a function of late summer rainfall and (b) decrease as a function of late summer temperature (*MR_final_*, Table 1). Some points offset in x and y planes to show the raw data; mean daily rainfall ranges between −1.76 and 2.18 (1.29–2.87 mm), and temperature ranges between −1.16 and 1.85 (15.61–18.11 °C). The condition (c) and phenophase (d) indices of *Geranium* at site G1 increase as a function of soil moisture. Quadrats with higher soil moisture are more likely to contain high proportions of good condition new recruits, and low proportions of poor-condition, reproductive *Geranium.* Soil moisture ranges between −1.58 and 1.83 (14.6–34.4 %).

**Table 2.** Parameter estimates (with standard errors) for the final (_*final*_; selected, most parsimonious) and null (_N_) logistic regression models investigating drivers of (a) the condition of the youngest *Geranium* (M*C*), and (b) the probability of new recruitment of *Geranium* (M*R*) in each quadrat during September surveys (2016–2017). Also presented for comparison are the restricted models (M*X_R_*) used to check for class bias in M*X_final_*. All models bar the null contain a subset of fixed effects from: site (S, using sum contrasts: see SI-1.9), daily mean rainfall (R) or daily mean temperature (T) in the period between August 1^st^–September sampling date. Other terms and time periods were tested and detected in the final candidate set of models (Table S8 and S9). β_0_ is the intercept, which accounts for the mean of site effects in all but the null models, *k* is the number of parameters, *LL* is the log-likelihood of the model and ΔAIC is the ΔAIC relative to the model with the lowest AIC in each case (Table S8 and S9).

| Model | Model parameters |  |  |  |  |  |  |  |
| --- | --- | --- | --- | --- | --- | --- | --- | --- |
| | $\beta_0$ | $S_1$ | $S_2$ | $R$ | $T$ | $k$ | $LL$ | $\Delta AIC$ |
| **(a) Condition of youngest *G. dissectum*** |  |  |  |  |  |  |  |  |
| **MC<sub>final</sub>** | 2.053<br>(0.535) | 5.749<br>(0.985) | -1.590<br>(0.551) | 5.871<br>(0.891) | – | 4 | -61.17 | 0.13 |
| **MC<sub>N</sub>** | 0.545<br>(0.141) | – | – | – | – | 1 | -143.30 | 168.38 |
| **MC<sub>R</sub>** | 1.565<br>(0.622) | 5.071<br>(1.116) | -1.305<br>(0.630) | 5.611<br>(1.128) | – | 4 | * | * |
| **(b) Probability of *G. dissectum* recruitment** |  |  |  |  |  |  |  |  |
| **MR<sub>final</sub>** | 0.310<br>(0.354) | 4.366<br>(0.794) | -3.558<br>(0.718) | – | -3.872<br>(0.555) | 4 | -49.07 | 1.43 |
| **MR<sub>N</sub>** | 0.352<br>(0.138) | – | – | – | – | 1 | -147.78 | 192.84 |
| **MR<sub>R</sub>** | -0.044<br>(0.495) | 4.770<br>(1.126) | -4.120<br>(1.017) | – | -4.096<br>(0.792) | 4 | * | * |

We had low statistical power to detect relationships between weather and the condition and recruitment of *Erodium,* models for which are outlined in SI-1.12.5.

### Microclimate effects on Geraniaceae phenology and condition

To address question (d), we assessed Geranium phenology and condition across a range of microclimates at site G1 in September 2017. At this site, areas of moister soil (where plants are less likely to dry out) were associated with Geranium plants in better condition and earlier phenophases (Figure 4c,d; Table 3). Candidate models suggested such plants were also more prevalent in areas with warmer, moister microclimates (SI-1.12.6).

**Table 3.**
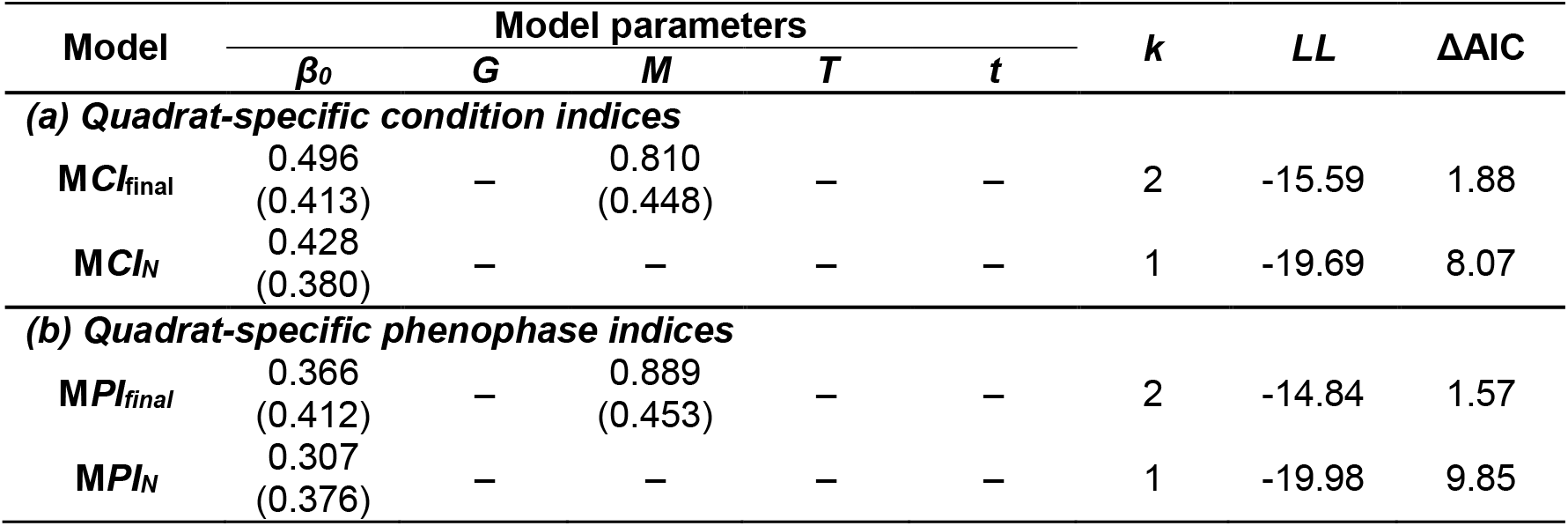
Parameter estimates (with standard errors) for the final (_*final*_; selected, most parsimonious) and null (_N_) logistic regression models investigating microclimatic drivers of (a) quadrat-specific condition indices (M*CI*), and (b) quadrat-specific phenophase indices (M*PI*) of *Geranium* at site G1 in September 2017. Also presented for comparison are the associated null models (M*X_N_*). All models bar M*X*_N_ contain a subset of fixed effects from: bare ground cover (G), soil moisture (M), mean (T) and minimum temperature (t). Other terms were tested and detected in the final candidate sets of models (Tables S9–10). β_0_ is the intercept, *k* is the number of parameters, *LL* is the log-likelihood of the model and ΔAIC is the ΔAIC relative to the model with the lowest AIC in each case (Tables S9–10).

| Model | Model parameters | | | | | $k$ | $LL$ | $\Delta AIC$ |
| --- | --- | --- | --- | --- | --- | --- | --- | --- |
| | $\beta_0$ | $G$ | $M$ | $T$ | $t$ | | | |
| *(a) Quadrat-specific condition indices* |  |  |  |  |  |  |  |  |
| $MCI_{final}$ | 0.496<br>(0.413) | — | 0.810<br>(0.448) | — | — | 2 | -15.59 | 1.88 |
| $MCI_N$ | 0.428<br>(0.380) | — | — | — | — | 1 | -19.69 | 8.07 |
| *(b) Quadrat-specific phenophase indices* |  |  |  |  |  |  |  |  |
| $MPI_{final}$ | 0.366<br>(0.412) | — | 0.889<br>(0.453) | — | — | 2 | -14.84 | 1.57 |
| $MPI_N$ | 0.307<br>(0.376) | — | — | — | — | 1 | -19.98 | 9.85 |

## Discussion

Here, we describe temporal variation in condition and phenology of three plant species and highlight the implications for their use as larval host plants by a butterfly species that has recently expanded its geographic range. Our data, from multiple sites across two years, suggest that the annual host *Geranium dissectum* varies more in condition and availability than both *Erodium cicutarium* and *Helianthemum nummularium*, the species that have been used as long-standing hosts at the range margin, and does so in a way that differs between years and with (micro-)climatic conditions. Such variation in condition and availability likely generates narrow and unpredictable phenological ‘windows of opportunity’ for exploitation of ephemeral annual species that vary among sites and years under the conditions of variable population sizes or phenology observed near the limits of species’ geographic ranges [37]. Though we recognise differences in the hosts other than their perennation strategy, in this case the perennial plant that was used as the main pre-expansion host appears to be a more reliable resource, where present, than the more widespread annual *Geranium* species that has acted as a primary host during the climate-associated range expansion. *Erodium,* is typically found on sandy soils, is relatively drought-tolerant compared to other Geraniaceae hosts, and appears to respond to a wider range of phenological cues than *Geranium,* which may improve its relative condition and availability as a host under the conditions we observed [30]. Our results suggest that climatic effects on the phenological synchrony of biotic interactions could act as a mechanism generating transient patterns of host associations at range margins, and consequently of habitat availability in the landscape and patterns of range shifting.

### (Micro-)climatic variation and host plant phenology

Our data indicate that the phenology of *Geranium,* a widespread annual host plant used by the brown argus butterfly, is sensitive to weather variability. Greenhouse experiments and field observations suggest that summer temperatures and moisture thresholds are crucial to dormancy breaking and germination in *Geranium* [38,39]. A complementary interpretation of these data is that *Geranium* plants may germinate early following a cool spell and early summer rain, but will suffer high seedling mortality where the summer is subsequently hot and dry (e.g. [40,41]). For example, our data show that hot, dry conditions in 2016 were associated with early and pronounced senescence of plants in July, as well as delayed germination and/or early seedling mortality. By contrast, our study sites received relatively high rainfall throughout summer 2017 [30], which is likely to have overcome moisture-dependency in dormancy breaking and/or promoted seedling survival. *Geranium* condition was also higher following wetter (2017) summer conditions, which supports evidence that drought and thermal stress cause premature senescence and declines in the quality (for consumers) of herbaceous plants [42,43]. A higher proportion of younger and better-condition *Erodium* were available in 2017 than in 2016, although we lacked statistical power to associate this with climatic variables (SI-1.12.5). However, our data support previous observations that reduced soil water availability advances the reproductive stage and the end of the growing season in *Erodium,* and reduce its investment in leaf biomass [44,45].

### Trophic interactions at range margins

Host condition and phenological synchrony in biotic interactions appear to be mediated by local (micro-)climatic variation, and are major determinants of spatiotemporal variation in fecundity and population size of host-specialist herbivores such as the brown argus [19,22,26,46,47]. By reducing temporal overlap between suitable resources and key herbivore life stages (e.g. adult egg-laying and early larval stages), adverse (micro-)climatic conditions may limit egg-laying and feeding opportunities and reduce larval survival, particularly where plants with limited temporal availability (such as the annual *Geraniaceae* studied here) are the main hosts [18,48]. Climatic conditions that are set to become more common (i.e. variable rainfall and longer, hotter summers [49,50]), may therefore narrow or close the phenological window of opportunity for this host-specialist herbivore to exploit these ephemeral annual resources in late summer [46,47].

Our analysis was unable to detect a clear geographic gradient in (a)synchrony, which may limit the predictability of the window of opportunity for brown argus to interact with Geraniaceae across its range. The widespread annual Geraniaceae are the hosts primarily used by brown argus at the expanding front of its recent range in Britain, whereas *Helianthemum* represents the apparent ancestral host at most sites where the species has been present for the past century or longer [26,30]. Therefore, while there is variation among sites, variation in butterfly-host synchrony (and the success of this relatively novel interaction) may be especially pronounced and unpredictable near the range limits, particularly if abiotic conditions are marginal and population sizes small [22]. In this respect, our results suggest that the range limit may be set not via broad environmental gradients in synchrony, but via shifting availability of sites where herbivore-host synchrony is sufficient. Through a process of ecological fitting at the novel range margin, populations interact with the resources that they happen to be synchronous with [51].

The relationships between weather, plant condition and phenology, which we identify here for *Geranium,* are therefore crucial in mediating how climate change, variability and unpredictability will affect synchrony in biotic interactions. Given that climate change is causing widespread changes in phenological synchrony, both existing and novel host interactions may be vulnerable under climate change. However, recent evidence suggests that at least as many biotic interactions are becoming more synchronous as are becoming less synchronous [22]. This evidence therefore also highlights the potential for novel biotic interactions to emerge, and further supports a role for spatiotemporal variation in synchrony underlying transient host interactions.

Spatiotemporal variation in the synchrony of biotic interactions is likely to generate transient mosaics of selection pressures for different diets, and thereby influence patterns of dietary specialisation/generalisation that recent research has shown to be an emergent and surprisingly common property of range dynamics [7,8,21,24]. In range expansions, many new consumer-resource interactions form and some may be lost [7,8]. For example, diet breadths in populations of Edith’s checkerspot butterfly *(Euphydryas editha)* increased after colonisation events as individual host preferences diversified, but populations subsequently reverted to monophagy [8].

Spatiotemporal variation in phenological synchrony may prove to be an underlying mechanism not only for lability in insect-host associations at range margins, but also for range limit stability under scenarios of temporal environmental variability. In particular, existing phenological plasticity can increase fitness costs where environments become more unpredictable [52], and range limits are more stable (expansions less likely) where environmental variance is too large for adaptation and colonisation [53]. Therefore, increasing environmental variance may preclude colonisation events that depend on predictably synchronous biotic interactions.

### Evolution during range expansion

Following the recent range expansion and incorporation of Geraniaceae into the diet of the brown argus, Geraniaceae-feeding populations that were able to persist became specialised on the novel hosts, losing the adaptive capacity to use *Helianthemum* [21,24]. Though specialisation on Geraniaceae appears to have become more reliable on average [29], our data suggest that interannual variation in phenology could still alter the success of this interaction, which may prove to be locally transient in the face of phenological asynchrony. In a comparable example from Scandinavia, narrow oviposition preferences of the Glanville fritillary *(Melitaea cinxia)* for phenologically-limited hosts risks high larval mortality under severe drought conditions in some years [20,54].

Conversely, phenological asynchrony at recently colonised Geraniaceae sites could lead to variable population dynamics, and selection favouring continued dispersal from the natal site in search of suitable egg-laying locations [55,56]. During the range expansion, selection has apparently favoured more dispersive phenotypes which have increased flight capacity and more readily accept the geographically widespread Geraniaceae hosts [24,27,28]. Consequently, under certain (micro-)climatic conditions, the dispersive, Geraniaceae-favouring phenotype may represent an alternative life history strategy that drives expansion at range margins and in-filling of the core range. Subsequent migrants that colonise *Helianthemum* sites may need to regain the ability to use *Helianthemum* (as shown in [21]; cf. [8]) in order to benefit from stability of (and phenological synchrony with) the host resource.

### Conservation and management implications

Understanding constraints and opportunities for species’ distributions is central to successful conservation practices. Our results highlight the importance of considering drivers of synchrony and the outcomes of biotic interactions when examining climate-driven range shifts, and recognising the crucial roles of microclimate and individual behaviour in mediating these interactions [19]. Conservation strategies could seek to maximise habitat and microclimatic heterogeneity to promote diversity in local phenologies across trophic levels [57,58]. In some cases, microclimatic variation may generate sufficient fine-scale spatial heterogeneity in relative phenology and host condition to buffer local herbivore populations against phenological asynchrony [10,18,59,60]. However, to improve our predictions of ecological responses to climate change, and of the critical levels of environmental change likely to cause rapid loss of ecosystem outputs, more empirical data are needed on shifts in biotic interactions across populations, climates and species ranges, their effects on demography, and their rates and patterns of evolution. In the present case, we have highlighted what might be typical variation in phenological synchrony across two years, which emphasises the potentially large indirect impacts of climate change on herbivore success. However, it would be beneficial to expand sampling across both time and space to better understand general patterns of interannual variation and gain a more holistic understanding of the phenomena discussed.

## Conclusions

Our results place a novel emphasis on the interactions between phenology, resource use and climate change in a range-expanding herbivore whose sensitivity to small changes in temperature might otherwise predict a positive range expansion response under future climates. Instead, small changes in temperature and moisture regime have the potential to disrupt the range expansion, mediated by phenological and physiological changes in the herbivore’s larval resource that essentially fragment the herbivore’s potential range. This suggests that novel host interactions may only remain as transient resources at shifting range margins. The mechanisms underlying insect-plant interactions, and their responses to climate change, are likely to be more complex than they appear, and we need more detailed knowledge of these mechanisms to understand and predict species’ interactions and responses to climate change.

## Supporting information

Erodium data

Site G1 Data

Summer Quadrat Data

All Quadrat Data

Geranium Data

Summary of supplementary material

## Acknowledgements

Research was funded by the Natural Environment Research Council (PhD studentship grant NE/L002434/1 to JES) and by the Botanical Society of Britain and Ireland (BSBI Science and Research Council grant to JES). Research permits were kindly granted by Butterfly Conservation, the Forestry Commission, Holkham Estate, the National Trust, Natural England, and Norfolk Wildlife Trust. Special thanks to R. Menéndez and J. Bennie for helpful comments on the research, and to R. Butlin, A. Phillimore and an anonymous reviewer whose comments greatly improved this work.

## References

1. Louthan AM, Doak DF, Angert AL. 2015 Where and when do species interactions set range limits? Trends in Ecology & Evolution 30, 780–792. (doi:10.1016/j.tree.2015.09.011)

2. Nadeau CP, Urban MC, Bridle JR. 2017 Coarse climate change projections for species living in a fine-scaled world. Global Change Biology 23, 12–24. (doi:10.1111/gcb.13475)

3. Sexton JP, McIntyre PJ, Angert AL, Rice KJ. 2009 Evolution and ecology of species range limits. Annual Review of Ecology, Evolution, and Systematics 40, 415–436. (doi:10.1146/annurev.ecolsys.110308.120317)

4. Macgregor CJ et al. 2019 Climate-induced phenology shifts linked to range expansions in species with multiple reproductive cycles per year. Nat Commun 10, 1–10. (doi:10.1038/s41467-019-12479-w)

5. Platts PJ, Mason SC, Palmer G, Hill JK, Oliver TH, Powney GD, Fox R, Thomas C. 2019 Habitat availability explains variation in climate-driven range shifts across multiple taxonomic groups. Scientific Reports 9, 15039.

6. Eriksson M, Rafajlović M. 2021 The role of phenotypic plasticity in the establishment of range margins. 2021.10.04.463099. (doi:10.1101/2021.10.04.463099)

7. Lancaster LT. 2020 Host use diversification during range shifts shapes global variation in Lepidopteran dietary breadth. Nature Ecology & Evolution 4, 963–969. (doi:10.1038/s41559-020-1199-1)

8. Singer MC, Parmesan C. 2021 Colonizations cause diversification of host preferences: A mechanism explaining increased generalization at range boundaries expanding under climate change. Global Change Biology 27, 3505–3518. (doi:10.1111/gcb.15656)

9. de la Paz Celorio-Mancera M, Wheat CW, Vogel H, Söderlind L, Janz N, Nylin S. 2013 Mechanisms of macroevolution: polyphagous plasticity in butterfly larvae revealed by RNA-Seq. Molecular Ecology 22, 4884–4895. (doi:10.1111/mec.12440)

10. Ward SF, Moon RD, Herms DA, Aukema BH. 2019 Determinants and consequences of plant–insect phenological synchrony for a non-native herbivore on a deciduous conifer: implications for invasion success. Oecologia 190, 867–878. (doi:10.1007/s00442-019-04465-2)

11. Visser ME, Holleman LJM, Gienapp P. 2006 Shifts in caterpillar biomass phenology due to climate change and its impact on the breeding biology of an insectivorous bird. Oecologia 147, 164–172. (doi:10.1007/s00442-005-0299-6)

12. Fuentealba A, Pureswaran D, Bauce É, Despland E. 2017 How does synchrony with host plant affect the performance of an outbreaking insect defoliator? Oecologia 184, 847–857. (doi:10.1007/s00442-017-3914-4)

13. Posledovich D, Toftegaard T, Wiklund C, Ehrlén J, Gotthard K. 2018 Phenological synchrony between a butterfly and its host plants: Experimental test of effects of spring temperature. Journal of Animal Ecology 87, 150–161. (doi:10.1111/1365-2656.12770)

14. Toftegaard T, Posledovich D, Navarro-Cano JA, Wiklund C, Gotthard K, Ehrlén J. 2019 Butterfly–host plant synchrony determines patterns of host use across years and regions. Oikos 128, 493–502. (doi:10.1111/oik.05720)

15. Schoonhoven LM, van Loon JJA, Dicke M. 2005 Insect-Plant Biology. Second Edition. Oxford, New York: Oxford University Press.

16. Kemp RJ, Hardy PB, Roy DB, Dennis RLH. 2008 The relative exploitation of annuals as larval host plants by European butterflies. Journal of Natural History 42, 1079–1093. (doi:10.1080/00222930801937608)

17. Ward SE, Schulze M, Roy B. 2018 A long-term perspective on microclimate and spring plant phenology in the Western Cascades. Ecosphere 9, e02451. (doi:10.1002/ecs2.2451)

18. Hellmann JJ, Weiss SB, McLaughlin JF, Ehrlich PR, Murphy DD, Launer AE. 2004 Structure and dynamics of *Euphydryas editha* populations. In On the Wings of Checkerspots. A Model System for Population Biology (eds PR Ehrlich, I Hanski), pp. 34–62. Oxford, United Kingdom: Oxford University Press.

19. Stewart JE, Maclean IMD, Edney EJ, Bridle JR, Wilson RJ. 2021 Microclimate and resource quality determine resource use in a range-expanding herbivore. Biology Letters 17, 20210175. (doi:10.1098/rsbl.2021.0175)

20. Rytteri S, Kuussaari M, Saastamoinen M. 2021 Microclimatic variability buffers butterfly populations against increased mortality caused by phenological asynchrony between larvae and their host plants. Oikos 130, 753–765. (doi:https://doi.org/10.1111/oik.07653)

21. Buckley J, Bridle JR. 2014 Loss of adaptive variation during evolutionary responses to climate change. Ecology Letters 17, 1316–1325. (doi:10.1111/ele.12340)

22. Kharouba HM, Ehrlén J, Gelman A, Bolmgren K, Allen JM, Travers SE, Wolkovich EM. 2018 Global shifts in the phenological synchrony of species interactions over recent decades. Proc Natl Acad Sci USA 115, 5211–5216. (doi:10.1073/pnas.1714511115)

23. Asher J, Warren MS, Fox R, Harding P, Jeffcoate G, Jeffcoate S. 2001 The Millennium Atlas of Butterflies In Britain and Ireland. Oxford, United Kingdom: Oxford University Press.

24. Bridle JR, Buckley J, Bodsworth EJ, Thomas CD. 2014 Evolution on the move: specialization on widespread resources associated with rapid range expansion in response to climate change. Proceedings of the Royal Society B: Biological Sciences 281, 20131800. (doi:10.1098/rspb.2013.1800)

25. Thomas CD, Bodsworth EJ, Wilson RJ, Simmons AD, Davies ZG, Musche M, Conradt L. 2001 Ecological and evolutionary processes at expanding range margins. Nature 411, 577–581. (doi:10.1038/35079066)

26. Bourn NAD, Thomas JA. 1993 The ecology and conservation of the brown argus butterfly *Aricia agestis* in Britain. Biological Conservation 63, 67–74. (doi:10.1016/0006-3207(93)90075-C)

27. Buckley J, Butlin RK, Bridle JR. 2012 Evidence for evolutionary change associated with the recent range expansion of the British butterfly, *Aricia agestis,* in response to climate change. Mol. Ecol. 21, 267–280. (doi:10.1111/j.1365-294X.2011.05388.x)

28. Bodsworth EJ. 2002 Dispersal and Behaviour of Butterflies in Response to Their Habitat. PhD thesis, University of Leeds, Leeds, United Kingdom.

29. Pateman RM. 2012 Climate Change and Habitat Associations at Species’ Range Boundaries. PhD thesis, University of York.

30. Stewart JE. 2020 Linking Phenology to Population Dynamics and Distribution Change in a Changing Climate. PhD Thesis, University of Exeter.

31. Suggitt AJ, Gillingham PK, Hill JK, Huntley B, Kunin WE, Roy DB, Thomas CD. 2011 Habitat microclimates drive fine-scale variation in extreme temperatures. Oikos 120, 1–8. (doi:10.1111/j.1600-0706.2010.18270.x)

32. Bramer I et al. 2018 Advances in monitoring and modelling climate at ecologically relevant scales. Next Generation Biomonitoring, part 1 58, 101–161. (doi:http://dx.doi.org/10.1016/bs.aecr.2017.12.005)

33. Dennis EB, Morgan BJT, Freeman SN, Roy DB, Brereton T. 2016 Dynamic models for longitudinal butterfly data. JABES 21, 1–21. (doi:10.1007/s13253-015-0216-3)

34. Met Office. 2017 UKCP09: Met Office gridded land surface climate observations - daily temperature and precipitation at 5km resolution. Updated to 2017.

35. Donoso I, Stefanescu C, Martinez-Abrain A, Treveset A. 2016 Phenological asynchrony in plant-butterfly interactions associated with climate: a community-wide perspective. Oikos 125, 1434–1444.

36. Harrison XA, Donaldson L, Correa-Cano ME, Evans J, Fisher DN, Goodwin CED, Robinson BS, Hodgson DJ, Inger R. 2018 A brief introduction to mixed effects modelling and multi-model inference in ecology. PeerJ 6, e4794. (doi:10.7717/peerj.4794)

37. Oliver TH, Roy DB, Brereton T, Thomas JA. 2012 Reduced variability in range-edge butterfly populations over three decades of climate warming. Global Change Biology 18, 1531–1539. (doi:10.1111/j.1365-2486.2012.02659.x)

38. Caudle C, Baskin JM. 1968 The germination pattern of three winter annuals. Bulletin of the Torrey Botanical Club 95, 331–335. (doi:10.2307/2483867)

39. Gama-Arachchige NS, Baskin JM, Geneve RL, Baskin CC. 2012 The autumn effect: timing of physical dormancy break in seeds of two winter annual species of Geraniaceae by a stepwise process. Ann Bot 110, 637–651. (doi:10.1093/aob/mcs122)

40. Baskin JM, Baskin CC. 1971 Germination of winter annuals in July and survival of the seedlings. Bulletin of the Torrey Botanical Club 98, 272–276. (doi:10.2307/2483627)

41. D’Aguillo MC, Edwards BR, Donohue K. 2019 Can the environment have a genetic basis? A case study of seedling establishment in *Arabidopsis thaliana*. J Hered 110, 467–478. (doi:10.1093/jhered/esz019)

42. Sade N, del Mar Rubio-Wilhelmi M, Umnajkitikorn K, Blumwald E. 2018 Stress-induced senescence and plant tolerance to abiotic stress. Journal of Experimental Botany 69, 845–853. (doi:10.1093/jxb/erx235)

43. Chaves MM, Maroco J, Pereira JS. 2003 Understanding plant responses to drought - from genes to the whole plant. Functional Plant Biology 30, 239–264.

44. Fresnillo Fedorenko DE, Fernández OA, Busso CA, Elia OE. 1996 Phenology of *Medicago minima* and *Erodium cicutarium* in semi-arid Argentina. Journal of Arid Environments 33, 409–416. (doi:10.1006/jare.1996.0076)

45. Cox JA, Conran JG. 1996 The effect of water stress on the life cycles of *Erodium crinitum* and *Erodium cicutarium* (Geraniaceae). Australian Journal of Ecology 21, 235–240. (doi:10.1111/j.1442-9993.1996.tb00604.x)

46. Abarca M, Spahn R. 2021 Direct and indirect effects of altered temperature regimes and phenological mismatches on insect populations. Current Opinion in Insect Science (doi:10.1016/j.cois.2021.04.008)

47. Hamann E, Blevins C, Franks SJ, Jameel MI, Anderson JT. 2021 Climate change alters plant–herbivore interactions. New Phytologist 229, 1894–1910. (doi:https://doi.org/10.1111/nph.17036)

48. Kemp R. 1998 Importance of larval foodplant lifespan to British butterflies - with particular reference to the Brown Argus *(Aricia agestis)*. Bulletin of the Amateur Entomologists’ Society 57, 225–227.

49. IPCC. 2014 Climate Change 2014: Synthesis Report. Contribution of Working Groups I, II and III to the Fifth Assessment Report of the Intergovernmental Panel on Climate Change (Core Writing Team, R.K. Pachauri and L.A. Meyer (eds.)). 151.

50. Wang J, Guan Y, Wu L, Guan X, Cai W, Huang J, Dong W, Zhang B. 2021 Changing lengths of the four seasons by global warming. Geophysical Research Letters 48, e2020GL091753. (doi:https://doi.org/10.1029/2020GL091753)

51. Janzen DH. 1985 On Ecological Fitting. Oikos 45, 308–310. (doi:10.2307/3565565)

52. Hoffmann AA, Bridle J. 2021 The dangers of irreversibility in an age of increased uncertainty: revisiting plasticity in invertebrates. Oikos (doi:10.1111/oik.08715)

53. Benning JW, Hufbauer RA, Weiss-Lehman C. 2021 Increasing temporal variance leads to stable species range limits. 2021.08.09.455156. (doi:10.1101/2021.08.09.455156)

54. Salgado AL, DiLeo MF, Saastamoinen M. 2020 Narrow oviposition preference of an insect herbivore risks survival under conditions of severe drought. Functional Ecology 34, 1358–1369. (doi:10.1111/1365-2435.13587)

55. Saastamoinen M et al. 2018 Genetics of dispersal. Biol Rev Camb Philos Soc 93, 574–599. (doi:10.1111/brv.12356)

56. Hanski IA. 2011 Eco-evolutionary spatial dynamics in the Glanville fritillary butterfly. PNAS 108, 14397–14404. (doi:10.1073/pnas.1110020108)

57. Suggitt AJ et al. 2018 Extinction risk from climate change is reduced by microclimatic buffering. Nature Clim Change 8, 713–717. (doi:10.1038/s41558-018-0231-9)

58. Oliver T, Roy DB, Hill JK, Brereton T, Thomas CD. 2010 Heterogeneous landscapes promote population stability. Ecology Letters 13, 473–484. (doi:10.1111/j.1461-0248.2010.01441.x)

59. Connor EF, Adams-Manson RH, Carr TG, Beck MW. 1994 The effects of host plant phenology on the demography and population dynamics of the leaf-mining moth, *Cameraria hamadryadella* (Lepidoptera: Gracillariidae). Ecological Entomology 19, 111–120. (doi:10.1111/j.1365-2311.1994.tb00400.x)

60. Bennett NL, Severns PM, Parmesan C, Singer MC. 2015 Geographic mosaics of phenology, host preference, adult size and microhabitat choice predict butterfly resilience to climate warming. Oikos 124, 41–53. (doi:10.1111/oik.01490)

