## Supplementary material for "Climate-driven variation in biotic interactions provides a narrow and variable window of opportunity for an insect herbivore at its ecological margin": SI-1

**<u>Supplementary Information 1. Supplementary methods and results</u>**

Appendix 1 to '*Climate-driven variation in biotic interactions provides a narrow and variable window of opportunity for an insect herbivore at its ecological margin'.* James E. Stewart, Ilya M.D. Maclean, Gara Trujillo, Jon Bridle and Robert J. Wilson.

### SI-1.1. Survey dates

We surveyed ten sites fortnightly–monthly between July 2016 and October 2017, to monitor phenology and condition of three host plant species of the brown argus butterfly, *Aricia agestis*. Site information (Table S1), survey information (Table S2) and site profiles are outlined below.

Table S1. Location (latitude and longitude), dominant and (subsidiary) host plants, daily mean summer temperature (°C) and daily mean summer rainfall (mm) in 2016 and 2017 for all study sites, arranged by descending latitude (north to south). Temperature and rainfall data are based on daily 5km gridded observations from the UK Met Office (Met Office, 2017). Summer is defined here as July 1st until the mean date of the early September plant surveys across sites and years (September 6th). E = *Erodium cicutarium*, G = *Geranium dissectum*, Gm = *G. molle*, Gp = *G. pratense*, H = *Helianthemum nummularium*.

| Site Code | Lat. (°N) | Long. (°E) | Host | Daily mean rainfall (mm) | | Daily mean temperature (°C) | |
|---|---|---|---|---|---|---|---|
| | | | | 2016 | 2017 | 2016 | 2017 |
| E1 | 52.975 | 0.779 | E (Gm) | 1.25 | 1.84 | 17.51 | 16.66 |
| E2 | 52.974 | 0.545 | E (Gm) | 1.50 | 2.11 | 17.67 | 16.89 |
| H1 | 52.630 | -0.415 | R | 1.19 | 1.76 | 17.86 | 16.91 |
| G1 | 52.343 | -1.454 | G | 1.19 | 2.04 | 17.21 | 16.31 |
| H2 | 52.231 | 0.363 | H | 0.93 | 2.29 | 18.23 | 17.18 |
| H3 | 51.858 | -0.551 | H | 1.01 | 2.54 | 17.39 | 16.41 |
| G2 | 51.671 | -1.266 | G (Gp, Gm) | 0.78 | 2.09 | 18.10 | 17.25 |
| H4 | 51.618 | -1.034 | H | 2.47 | 2.47 | 17.11 | 16.27 |
| H5 | 51.059 | -1.280 | H | 1.48 | 4.28 | 17.15 | 16.46 |
| G3 | 51.058 | -1.275 | G (H) | 1.48 | 4.28 | 17.15 | 16.46 |

Table S2. Survey dates (DD/MM/YY format) for all Geraniaceae sites surveyed, including details of the number of quadrat locations [x] surveyed at each visit.

| Site | | | | | | | | | |
|---|---|---|---|---|---|---|---|---|---|
| **E1** | **E2** | **G1** | **G2** | **G3** | **H1** | **H2** | **H3** | **H4** | **H5** |
| 28/07/16 [30] | 27/07/16 [30] | 24/07/16 [30] | 23/07/16 [34] | 20/07/16 [30] | 26/07/16 [30] | 30/07/16 [30] | 31/07/16 [30] | 22/07/16 [30] | 21/07/16 [30] |
| 11/08/16 [32] | 10/08/16 [32] | 09/08/16 [30] | 07-10/08/16 [47] | 13/08/16 [30] | 06/09/16 [30] | 09/09/16 [30] | 10/09/16 [30] | 08/08/16 [20] | |
| 7/09/16 [30] | 07-08/09/16 [32] | 05/09/16 [30] | 12/09/16 [32] | 03-04/09/16 [30] | 08/10/16 [30] | 10/10/16 [30] | 11/10/16 [30] | 11/09/16 [30] | 16-18/09/16 [36] |
| 9/10/16 [30] | 09/10/16 [30] | 06/10/16 [40] | 12/10/16 [40] | 12/10/16 [35] | | | | 11/10/16 [30] | 13/10/16 [30] |
| - | - | - | 30/10/16 [30] | 31/10/16 [30] | | | | 02/11/16 [10] | 31/10/16 [10] |
| - | - | - | 02/12/16 [30] | 04-05/12/16 [30] | | | | 03/12/16 [20] | 05/12/16 [20] |
| - | - | - | 04/01/17 [30] | 05/01/17 [30] | | | | 04/01/17 [20] | 05/01/17 [20] |
| 6/02/17 [20] | 06/02/17 [20] | - | 05/02/17 [30] | 04/02/17 [25] | | | | 05/02/17 [15] | 04/02/17 [10] |
| 11/03/17 [25] | 10-11/03/17 [34] | - | 08/03/17 [26] | 07/03/17 [21] | | | | 08/03/17 [20] | 07/03/17 [20] |
| 07/04/17 [30] | 07/04/17 [30] | 13/04/17 [20] | 11/04/17 [41] | 09/04/17 [20] | | | | | 09/04/17 [20] |
| 14/05/17 [27] | 13/05/17 [37] | 17/05/17 [60] | 18/05/17 [60] | 11/05/17 [30] | | | | | |
| 02/06/17 [39] | 03/06/17 [38] | 07/06/17 [60] | 07/06/17 [60] | 08/06/17 [30] | | | | | |
| 23-24/06/17 [47] | 25/06/17 [39] | 28/06/17 [60] | 29/06/17 [60] | 30/06/17 [31] | | | | | |
| 30/07/17 [46] | 29/07/17 [39] | 01/08/17 [60] | 27/07/17 [59] | 02/08/17 [35] | | | | | |
| 15/08/17 [46] | 14/08/17 [39] | 17-18/08/17 [60] | 12/08/17 [59] | 18/08/17 [30] | | | | | |
| 30/08/17 [46] | 31/08/17 [38] | 06/09/17 [60] | 29/08/17 [55] | 07/09/17 [30] | | | | | |
| 10/10/17 [40] | 09/10/17 [35] | 12/10/17 [58] | 07/10/17 [30] | 03/10/17 [30] | | | | | |

### SI-1.2. Site profiles

#### Geraniaceae sites

*Site E1: Holkham National Nature Reserve*

Holkham National Nature Reserve (52.975 °N 0.779 °E, Ordnance Survey grid reference TF 870 450; site E1 in Figure 1, main text) lies on the north Norfolk coast, east of Holme Dunes NNR (site E2). The reserve is composed of a variety of habitat types, including salt and grazing marshes, woodland, sand dunes and foreshore. *E. cicutarium* is common, and *G. molle* occasional, in the sand dune areas, particularly where rabbits are active. This coastal dune grassland is the primary habitat for *Aricia agestis* at E1. *G. dissectum* is uncommon and restricted to path-side verges and areas of more shaded, stable soil; *Helianthemum nummularium* is not known to occur at E1 (JES, *pers. obs.*).

*A. agestis* has been observed at E1 since the introduction of a butterfly transect to the site in 1976, as part of the United Kingdom Butterfly Monitoring Scheme (UKBMS) [1,2], and now forms a stable population within the pre-expansion range.

*Site E2: Holme Dunes National Nature Reserve*

Holme Dunes National Nature Reserve (52.974 °N 0.545 °E, OS grid reference TF 714 450; site E2 in Figure 1, main text) lies on the north Norfolk coast and is composed of a range of coastal habitats from the intertidal sand and mudflats to saltmarsh, grazing marsh, sand dunes and limited coniferous woodland. *E. cicutarium* is common, and *G. molle* occasional, in the sand dunes and rabbit-grazed grassland areas, though *G. dissectum* is scarce and typically restricted to the grazing meadows (JES, *pers. obs.*). *H. nummularium* is absent from the site, and the nearest known record is from Ringstead Down (Ordnance Survey grid reference TF 704 401, 4.9km ESE of E2; (NBN Atlas, 2017)). Egg-laying by *A. agestis* has been observed on both *E. cicutarium* and *G. molle* in the dune system (JES, *pers. obs*).

A UKBMS transect was established at E2 in 1978, and *A. agestis* has been recorded there since 1985 [1,2], though it is known to have been present in the area from before this, and the site is within the pre-expansion range of *A. agestis* [3].

*Site G1: Bubbenhall Meadow*

Bubbenhall Meadow (52.343 °N 1.454 °W, OS grid reference SP 373 717; site G1 in Figure 1, main text) is an area of neutral grassland in Warwickshire. G1 is bordered by mixed woodland and has recently been widely planted with young saplings across much of the meadow in order to form a woodland corridor. By the end of the plant surveys in 2017, tree growth across G1 was still minimal, resulting in negligible shading of the grassland. Topographically, G1 resembles a large, lightly undulating bowl and therefore has a variety of topographic microclimates formed by, for example, south-facing and east-facing slopes. Some small patches of scrub exist, which further contribute to shading and microclimatic variation. *G. dissectum* is relatively common within the meadow, though the other host plants have not been observed at the site (JES, *pers. obs.*).

A UKBMS butterfly transect was established at G1 in 2008, and *A. agestis* has been observed on-transect since then [1,2]; however, there are not enough data to classify *A. agestis'* population status at the site. The pre-expansion range of *A. agestis* does not include G1, though it is known to have been present in the same 10 km square since at least as early as 1999.

*Site G2: Barton Fields*

Barton Fields (51.618 °N 1.266 °W, OS grid reference SU 507 972; site G2 in Figure 1, main text) is an area of grassland, scrub and wet meadow on the eastern outskirts of Abingdon, Oxfordshire. G3 is managed for wildlife conservation by the local naturalists' society such that some areas of the grassland are mown annually in early autumn, but others are left to grow. *G. dissectum* is present throughout the grassland area but is less frequent in the western end. The western end of the grassland is in close proximity to the wet meadow area and infrequently experiences minor flooding. In this western end, *G. pratense* is more common. *G. molle* is rare, whilst *E. cicutarium* and *H. nummularium* are not known at G3.

*A. agestis* has been observed at G3 since the UKBMS butterfly transect began there in 2011 [1,2]. Due to the site's location within the pre-expansion range of *A. agestis*, it is likely that *A. agestis* used G3 before 2011.

*Site G3: Magdalen Hill Down 'Extension'*

Magdalen Hill Down 'Extension' (51.058 °N 1.275 °W, OS grid reference SU 515 292; site G3 in Figure 1, main text) is so-called because it is an extension of the Butterfly Conservation reserve of Magdalen Hill Down, east of Winchester in Hampshire. G3 was an area of calcareous grassland transformed for intensive arable cultivation; it was in arable use for over 50 years until 1995, when it was purchased by Butterfly Conservation and rehabilitated to flower-rich grassland with limited scrub areas at the borders. The western edge of G3 is only separated from R5 by a fence and a line of trees and scrub, but supports markedly different vegetation, likely due to the recent history of agricultural inputs to the soil. The characteristic chalk downland flora is gradually being recreated through sowing of native grasses and wildflower species from nearby sites. *G. dissectum* is frequent in the field margins and in some of the grassland areas, most noticeably in the southwest of the site. A similar species, *G. columbinum* (long-stalked crane's-bill), is also present in smaller numbers, while *G. molle* is rare on-site. *E. cicutarium* is not known at the site, and *H. nummularium* is restricted to a small number of artificial chalk scrapes (JES, *pers. obs.*). Evidence of egg-laying has been observed on both *G. dissectum* and *H. nummularium* at G3 (JES, *pers. obs.*).

UKBMS transects began at G3 in 1996, and *A. agestis* has been seen since the first year of transects [1,2]. However, *A. agestis* is known to have been observed in the area prior to 1990: sightings were recorded in the relevant 10 km grid square during the first national butterfly atlas, spanning the period 1970–1982 [3]. The UKBMS index at G3 suggests that *A. agestis* is in decline, and it is unclear whether individuals move between G3 and the neighbouring *H. nummularium* site (H5, Figure 1, main text), or indeed whether H5 acts as a source or sink for this population. Evidence of egg-laying has been observed on both *G. dissectum* and *H. nummularium* at G3 (JES, *pers. obs.*).

**Helianthemum nummularium sites**

*Site H1: Barnack Hills and Holes NNR*

Barnack Hills and Holes National Nature Reserve (52.630 °N 1.415 °W, OS grid reference TF 070 040; site H1 in Figure 1, main text) is a topographically diverse area of limestone grassland and mixed scrub in the north-west of Cambridgeshire. *H. nummularium* is the dominant host here, and is very abundant across the grassland areas [4] (JES, *pers. obs.*). *G. molle* and *G. dissectum* both occur at the site, but are very rare and typically restricted to small areas of scrub and path edges. *E. cicutarium* is not known to occur at H1.

A UKBMS transect was established at the site in 1981 but *A. agestis* was not recorded at the site until 1994 [1,2]. However, *A. agestis* is thought to have been present at the site before 1981 (C. Gardiner, *pers. comm.*) and H1 is within the pre-expansion range. Natural egg-laying has only been observed on *H. nummularium* at H1 (C. Gardiner, *pers. comm.*), though extensive searches have not been performed on the Geraniaceae. Experimental work has demonstrated an ability of brown argus to lay on Geraniaceae hosts at this site [5].

*Site H2: Devil's Dyke SSSI*

Devil's Dyke (52.231 °N 0.363 °E, OS grid reference TL 615 617; site H2 in Figure 1, main text) is a Site of Special Scientific Interest in eastern Cambridgeshire. It is a linear earth bank of Anglo-Saxon origin that stretches for 12km from the north-west to south-east, though the present study focussed on a 1.5km stretch adjacent to Newmarket Racecourse. H2 is dominated by open calcareous grassland, with occasional scrub patches. *H. nummularium* is very abundant along the bank, particularly along the south-west-facing bank. The Geraniaceae hosts are not known to occur at the site except in small shaded patches near to tree growth where *G. dissectum* is rarely found.

H2 is within the pre-expansion range of *A. agestis*, which is known to have been present the site since at least 2003, when a UKBMS transect was established at H2 [1,2].

*Site H3: Whipsnade Downs*

Whipsnade Downs (51.858 °N 0.551 °W, OS grid reference TL 012 182; site H3 in Figure 1, main text) is an area of calcareous grassland in Bedfordshire. *H. nummularium* is the dominant host plant at the site, though *G. dissectum* occurs rarely in ranker grassland areas, particularly near disturbed ground and in areas sheltered by the limited available tree cover (JES, *pers. obs.*).

A UKBMS transect was established at H3 in 1988. *A. agestis* has been observed on-site during all years in which the transect has been monitored to date [1,2]. H3 is within the core stronghold of *A. agestis*' distribution; individuals are known to have been recorded within the relevant 10 km grid square since at least 1980, prior to the range expansion.

*Site H4: Swyncombe Downs*

Swyncombe Downs (51.618 °N 1.034 °W, OS grid reference SU 668 915; site H4 in Figure 1, main text) is an area of south-facing calcareous grassland in Oxfordshire. The habitat in this SSSI is typical of chalk downland in the Chilterns, dominated by open grassland with small patches of mixed woodland. *H. nummularium* is very common at H4, and is thought to be the only host plant present on-site (JES, *pers. obs*.). Experimental work has demonstrated an ability of brown argus to lay on Geraniaceae hosts at this site [5].

*A. agestis* has been observed at H4 since monitoring began in 2002 as part of the UKBMS [1,2]. It has also been seen at a nearby site (Aston Rowant NNR: 51.667 °N 0.953 °W, OS grid reference SU 725 970) since UKBMS monitoring began in 1976 [2]. The area, part of an extensive stretch of calcareous grassland, is thought to have been a stronghold for *A. agestis* during the range retraction and is now within the core of its UK distribution [3,6].

*Site H5: Magdalen Hill Down 'Original'*

Magdalen Hill Down is a large Butterfly Conservation Reserve due east of Winchester in Hampshire (51.059 °N 1.280 °W, OS grid reference SU 505 920; site H5 in Figure 1, main text). Magdalen Hill Down 'Original' is so-called as it is the original section of calcareous downland in the area reclaimed by Butterfly Conservation for nature conservation purposes. Prior to reclamation in 1989, the downland was dominated by scrub but is now an open area of calcareous grassland that supports a range of species typical to this habitat type, including common bird's-foot-trefoil (*Lotus corniculatus*), thyme (*Thymus spp.*), marjoram (*Origanum vulgare*) and *H. nummularium*, which is very abundant on the south-facing slopes of the reserve. The Geraniaceae have not been observed on this section of the Magdalen Hill Down reserve, though *G. dissectum* is relatively common in some areas of the reserve's "Extension" (site G3).

A UKBMS transect was established at the site in 1990, and *A. agestis* has been recorded there in every year since then [1,2]. *A. agestis* is also known to have been observed in the area (part of the pre-expansion range) prior to 1990: sightings were recorded in the relevant 10 km grid square during the first national butterfly atlas, spanning the period 1970–1982 [3].

### SI-1.3 Quadrat surveys

*SI-1.3.1 Quadrat placement*

Key plant characteristics relevant to phenology and condition (outlined in the Main Text) were measured in 50 × 50 cm (0.25 $m^2$) quadrats at each of ten sites between July 2016 – October 2017. On average we assessed 33.7 quadrat samples per visit at each site. Quadrats were placed pseudo-randomly: random points were generated using ArcGIS (ArcGIS 10.2.1 for Desktop, ESRI, California, USA) for each site, typically along the approximate route of the UKBMS butterfly transect or in accessible locations. These points were located *in situ* using a handheld GPS (GPSmap 60CSx, Garmin, Kansas, USA). Quadrats were then placed over the patch of host plant that was nearest to the random point. Quadrats were thrown haphazardly to the ground to avoid biased selection of vegetation characteristics within a patch of host plants (for example, selection for plant condition or ground cover). The number of quadrats surveyed at each visit is summarised in Table S2. Minimum convex polygons encompassing all quadrats at each site had an average area of 1.69 ha +/- 1.79 ha.

*SI-1.3.2 Assessing vegetation characteristics*

For each host plant species found within the 50 × 50 cm quadrat, we recorded host plant percentage cover (below), phenophase (phenological stage) and condition. We also measured to attributes known to affect fine-scale microclimate: mean sward height (based on sward height in each corner of the quadrat) and percentage cover of bare ground [7,8]. String was used to divide the quadrat into 25 equally-sized (4%) sections which were used to estimate percentage cover of plants and bare ground to the nearest 0.5 %.

Condition (*Q*) was visually assessed on a scale of 0–3, based on plant colour, senescence/ wilting and shine (Figure S1), following [9,10]. Host condition, defined in this way, appears to align with the quality of the host material for the brown argus, which choose good condition plants over poor condition plants for egg-laying [9,10]. See ‘Assessing host condition’, below. For the Geraniaceae, category 0 includes plants that appear dead (browned and brittle vegetative tissue); category 1 plants are entirely or almost entirely composed of dull, yellowed and/ or wilted tissue and which may have some dead parts or dull green tissue; category 2 includes plants that are still mostly a dull green and which may have begun to show signs of wilting; category 3 plants have lush green leaves with little signs of senescence. *Helianthemum nummularium* follows similar criteria although, as established by Bourn and Thomas [9], category 2 *H. nummularium* may also present shiny yellow leaves, while category 3 *H. nummularium* leaves are green and shiny [9]. Phenophase was recorded on a four-point scale describing whether the plant was in leaf, bud, flower or had set seed.

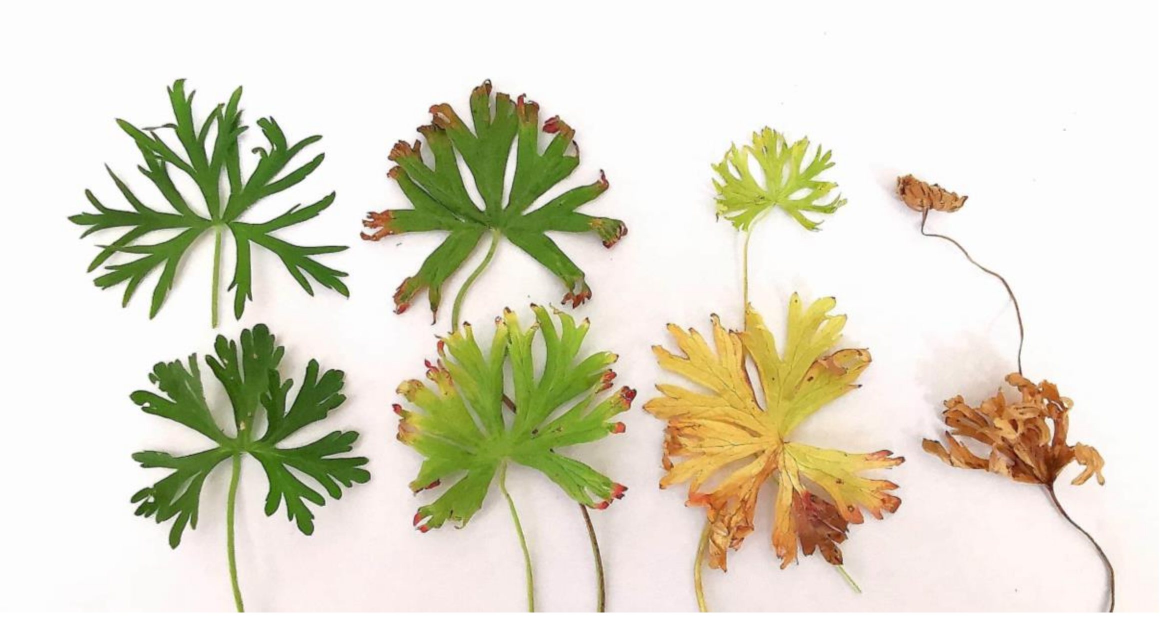

Figure S1. Leaf specimens from wild-grown *Geranium dissectum*, indicating differences in plant condition on a scale from 0 (poor condition; far right) to 3 (best condition; far left).

*SI-1.3.3 Assessing host condition*

We defined host condition based on visual assessments of the degree of desiccation/senescence and ‘greenness’ of the plant tissue, recognising that herbivore host selection is a complex process that is likely to be driven by more than variation in visually-perceptible traits. However, a range of related evidence provides confidence in our application of a visual assessment of resource condition, outlined below.

The ‘greenness’ of plants is a commonly applied proxy for productivity, for example in visual assessments and satellite-derived indices based on normalised difference vegetation index (NDVI) [9–12]. More specifically relevant to this study is that our plant condition scale was based on that applied by Bourn and Thomas [9], who demonstrated that, when laying on *H. nummularium*, the brown argus prefers to lay its eggs on “lush” shiny green leaves  (category 3). Bourn and Thomas [9] showed that this visual index of resource condition was associated with increased leaf mesophyll thickness and nitrogen content. For the Geraniaceae hosts our visual condition categories appear to correlate well with the egg-laying preferences of adult brown argus. Experimental work has demonstrated that the plant condition scale correlates with brown argus egg-laying behaviour on the Geraniaceae hosts: adult females lay significantly more eggs on good condition (‘high quality’) plants than on poor condition plants [10].

### SI-1.4 Quantifying variation in host condition and phenology

To test our expectation for greater temporal variation in annual vs perennial host condition we used a Kruskal-Wallis test for each sampling period to compare quadrat-level condition between host species. Bonferroni-corrected Dunn's tests were then used to identify which hosts differed significantly from one another in condition score within each sampling period. Quadrats were able to be treated as independent for each comparison, but this analysis was unable to account for among-site variation in host condition within each comparison. However, the among-site variation is shown in Figure 2a.

We then conducted interannual comparisons of Geraniaceae host condition (Mann-Whitney U tests) and phenophase ($X^2$ tests): these compared 2016 data with 2017 data, separately for each month between July to October. These months are the most relevant for host choice and larval feeding by second generation brown argus and their offspring. Quadrats were placed in different locations in 2016 compared to 2017, so could be considered independent between years.

### SI-1.5 Calculating host condition and phenology indices

#### *SI-1.5.1 Site-level indices*

We calculated host condition indices based on quadrat survey data for all summer survey dates (July – October 2016 and 2017); these indices were then used to estimate synchrony with brown argus adult and larval stages.

The site-level condition index is based on a mean of three quotient terms. The three terms are the antecedent terms in each of three quotients, *A-C*, expressed as ratios. For each site, *A* is the number of quadrats assigned the condition category 'c3' at each survey divided by the sum of quadrats of all categories (c3, c2, c1, and c0). *B* is the sum of 'c3' and 'c2' quadrats at each survey divided by the sum of quadrats of all condition categories. *C* is the sum of condition 'c3', 'c2', and 'c1' quadrats at each survey divided by the sum of all quadrat condition categories. Summing the antecedent terms of these three ratios gives *D*. *D*, when divided by *r* (the number of terms, here three), gives a condition index which ranges from zero (all in worst condition) to one (all in best condition). That is, the site-level condition index, $P_S = D/r$, where

$$D = A + B + C, \text{ and}$$

$$A = c3 / (c3 + c2 + c1 + c0),$$

$$B = c3 + c2 / (c3 + c2 + c1 + c0),$$

$$C = c3 + c2 + c1 / (c3 + c2 + c1 + c0),$$

and $r$ = 3.

Take for example a site with 20 c3, 5 c2, 2 c1 and 3 c0 quadrats (site 1) and a site (site 2) with 8 c3, 10 c2, 8 c1 and 4 c0 quadrats. The condition index for each site ($Q_{S1}$ *and* $Q_{S2}$) are calculated as follows:

$A_1 = c3_1 / (c3_1 + c2_1 + c1_1 + c0_1)$: $A_1 = 20 / (20 + 5 + 2 + 3) = 0.6667$

$B_1 = c3_1 + c2_1 / (c3_1 + c2_1 + c1_1 + c0_1)$: $B_1 = 25 / (30) = 0.8333$

$C_1 = c3_1 + c2_1 + c1_1 / (c3_1 + c2_1 + c1_1 + c0_1)$: $C_1 = 27 / (30) = 0.9$ ;

$D_1 = A_1 + B_1 + C_1$: $D_1 = 0.6667 + 0.8333 + 0.9 = 2.3999$

$\underline{P_{S1} = D_1/r: P_{s\,1} = 2.3999/3 = 0.7999}$

$A_2 = c3_2 / (c3_2 + c2_2 + c1_2 + c0_2)$: $A_2 = 8 / (8 + 10 + 8 + 4) = 0.2667$,

$B_2 = c3_2 + c2_2 / (c3_2 + c2_2 + c1_2 + c0_2)$: $B_2 = 18 / (30) = 0.6$,

$C_2 = c3_2 + c2_2 + c1_2 / (c3_2 + c2_2 + c1_2 + c0_2)$: $C_2 = 26 / (30) = 0.8667$,

$D_2 = A_2 + B_2 + C_2$: $D_2 = 0.2667 + 0.6 + 0.8667 = 1.7334$

$\underline{P_{S2} = D_2/r: P_{S2} = 1.7334/3 = 0.5778.}$

Geraniaceae host condition indices were calculated separately for each host species and survey date using data on the youngest plant in each quadrat. This approach is more informative for the Geraniaceae hosts than for *H. nummularium*, because *H. nummularium* host condition indices were only calculated for 2016, and used quadrat-level average condition classifications: finer-scale data on individual plants were not available, and *H. nummularium* sites were not surveyed in summer 2017.

*SI-1.5.2 Fine-scale condition and phenophase indices*

We calculated a quadrat-level condition index which accounted for the condition of all plants present within each quadrat at site G1. As above, the quadrat-level condition index is based on a mean of three quotient terms, and is calculated in the same way as the site-level index. The three terms are the antecedent terms in each of three quotients, *A-C*, expressed as ratios. For each quadrat, *A* is the number of plants assigned the condition category c3 at each survey divided by the sum of plants of all categories (c3, c2, c1 and c0). *B* is the sum of c3 and c2 plants at each survey divided by the sum of plants of all condition categories. *C* is the sum of c3, c2 and c1 plants at each survey divided by the sum of all plant categories. Summing the antecedent terms of these three ratios gives *D*. *D*, when divided by *r* (the number of terms, here three), gives a condition index which ranges from zero (all in worst condition) to one (all in best condition).

We also calculated a quadrat-level index of host phenophase, $P_Q$, by substituting our estimates by substituting our estimates of host plant phenophase into the equations above. By substituting category L for host condition c3, B for c2, F for c1 and S for c0, we calculated $P_Q$ such that it ranges between zero (all at the seed, or S, stage) to one (all in leaf, L, with no reproductive structures).

### SI-1.6 Brown argus phenology curves

We plotted values of the site-level host indices (above) against day of the year for all summer surveys (July–October), and overlaid these plots with Gaussian curves of the emergence phenology of adult second generation brown argus and their offspring. The phenology curves of second generation brown argus adults were calculated from output of a phenomenological model of phenology developed by Dennis *et al.* [13] and developed to account for effects of latitude and temperature by Stewart [14]. The phenomenological model was applied to counts of brown argus from 293 sites surveyed by the UK Butterfly Monitoring Scheme (UKBMS; [15]) between 1992–2016. These sites represent a subset of those surveyed by the UKBMS and excludes sites at which the focal species was not found, or at which monitoring was undertaken for fewer than five years.

We used the mean and standard deviation of the brown argus' peak flight date for the 1992–2016 period ([14]; *unpubl. data*) to plot the Gaussian phenology curves of second generation brown argus adults. We present a phenology curve for each site and year, accounting for variation due to site latitude and temperature, as outlined below. Output of the phenomenological model demonstrated that the absolute timing of second brood flight is necessarily linked to that of the first brood, which itself is negatively associated with latitude: the first brood emerges earlier further south (Table S3). Conversely, second brood phenology is earlier further north (as supported by recent studies [13,14]), but is also earlier under warmer summer conditions (Table S3). Specifically, second brood phenology is earlier when the 'between-brood temperature' is higher: the mean temperature (based on UK Met Office 5km gridded dataset for each site; [16]) between the peak of the first and second flight periods. The respective peaks used in calculation of the between brood temperature were calculated as the mean flight period of the first and second brood averaged across the 1992–2016 period, as reported by [14].

Table S3. Parameter estimates from phenomenological model, based on [14], showing estimates of the mean and standard deviation of the flight period for the first ($\mu_1$) and second brood ($\mu_2$) brown argus across 293 sites sampled by the UK Butterfly Monitoring Scheme (UKBMS). The exponent of $\mu_1$ is the mean flight date of the first brood in weeks from April 1st (annual start of UKBMS sampling). Therefore, $\exp(\mu_1) + \exp(\mu_2)$ is the mean flight date of the second brood. Also shown are the estimates for slopes of phenology on site northing and between-brood temperature (BBT). All covariates were standardised to zero mean and unit standard deviation, thus slopes represent comparable effect sizes.

| **$\mu_1$ parameter estimates** | | **$\mu_2$ parameter estimates** | | |
|---|---|---|---|---|
| **Intercept (mean ±SD)** | **Slope: north** | **Intercept (mean ±SD)** | **Slope: north** | **Slope: BBT** |
| 2.203 ± 0.757 | 0.036 | 2.424 ± 0.747 | -0.062 | -0.012 |

Using model-derived estimates ([14]; Table S3), we plotted site- and year-specific adult phenology curves in which the mean flight date was based on site latitude and the between-brood temperature in each year (2016 and 2017). The standard deviation of the flight date, used to determine the spread of the Gaussian curves, was also estimated by the model and was assumed to be consistent across sites and years. The Gaussian phenology curves of the offspring of these butterflies was based on the same mean flight dates and standard deviations, but shifted 11 days later to account for the time taken for mating and egg-laying (assumed to occur approximately 4 days after adult emergence) and egg development, which takes approximately one week [5]. There is likely to be variation around these timings (for example, based on local weather and population density); this means that the larval emergence curve is presented as an indicator rather than as a precise evaluation of appearance or abundance at a site.

### SI-1.7 Quantifying plant-herbivore asynchrony

The overlap between plotted condition indices and brown argus phenology curves was used to generate an area under the curve (AUC) metric of site- and year-specific synchrony between brown argus and each host plant. The AUC synchrony metric is derived from the plots in Figure 3 by taking the full area of each butterfly phenology curve, minus the area of the curve which lies above the corresponding host condition line for that site and year. The AUC metrics were then modelled in a beta regression (logit link) to test for effects of site latitude, host plant, year and a host-year interaction on estimated asynchrony. For each beta regression, n = 15, which limited the ability to successfully fit GLMs with maximum likelihood-based approaches, especially for models with an interaction term. Therefore, for adult and larval AUC metrics we fit Bayesian beta regressions with R package ‘brms’ [17], using vaguely informative normal priors for each of the regression parameters (defined in the attached R code). We used four Markov chains with 15000 iterations, including a 7500 iteration burn-in per chain. We performed model checks using R package ‘DHARMa’ [18], visual checks of chain mixing and model convergence, posterior predictive checks using R package ‘bayesplot [19], and used R package ‘performance’ [20] to calculate the conditional $R^2$ and perform leave-one-out cross-validation to derive Pareto k diagnostic estimates.

### SI-1.8 Choice of time periods for weather data in logistic regressions

We tested for climatic drivers of recruitment and condition of each Geraniaceae species in early September, when most larval offspring were expected to have emerged to feed. Using daily weather data [16], we calculated the minimum, mean and maximum temperature and mean rainfall for three periods in each year: July, July–September and August–September (only including weather data up to the day of quadrat sampling at each site in early September).

These three periods were chosen based on observations of declines in host condition and changes in host phenology, which suggested that where such changes occur, they may do so from late July through early September. Due to the seemingly rapid response of host plant condition to changes in temperature and rainfall, we consider weather data for July (the weeks preceding our late-July host plant surveys), July – September (summer weather over the key monitoring period up to our focal early-September surveys) and August-September (late summer weather for the approximately 5 weeks preceding our focal early-September surveys).

### SI-1.9 Model selection, validation and diagnostics

All models contained the term ‘site’ as a predictor, which was coded as ‘sum contrasts’. Under this formulation, the model intercept includes the mean of all site effects, $S_1$ shows the difference between the mean of level $S_1$ and the mean of all site effects, and $S_2$ shows the difference between the mean of level $S_2$ and the mean of all site effects.

For all logistic regressions, we constructed a complete model set based on all (reasonable) combinations of predictors. We reduced this full-subsets model set by removing models in which (1) estimated pairwise correlations between any two predictors was too high (Spearman’s $\rho > 0.5$), or variance inflation factors (using vif in ‘*car*’ [21]) indicated significant collinearity (VIF > 4); or (2) quadratic terms were present without their corresponding linear term. Model diagnostics were checked based on visual assessment of residual plots (residuals vs fitted values, QQ, scale-location and leverage plots), assessment of simulated residuals using the R package '*DHARMa*' [18] and deviance residuals with '*stats*' [22]. We then used AIC for model selection to create a *candidate set* of models.

The model having the lowest AIC value is likely to be the most parsimonious; however, as AIC is only an estimate of parsimony, we followed Richards [23] in considering certain other models as well. First, we highlighted a candidate set of models with AIC values within 6 units of the model with the lowest AIC value. In this *candidate set*, we used ΔAIC to denote the difference between the AIC value of model M and the lowest AIC value calculated. Thus, the best AIC model ($M_{best}$) had ΔAIC = 0 and all candidate models had a ΔAIC-value ≤ 6. Next, to prevent selecting unsupported, overly-complex models, we removed models from the candidate set that were more complex versions of other selected models [23]. Where this process failed to identify a single 'best' model, we follow Richards [24] in basing biological inference on the simplest (lowest *k*) model (or the best AIC model if *k* is equal across candidate models) and considering a parameter to have strong support if it is included in all candidate models. We denote the selected model $M_{final}$. This approach allows all good candidate models to be compared, and permits consideration of other important variables that would be excluded using methods such as stepwise selection [25].

The fit of the candidate set of logistic regression (binomial GLM) models was checked using the area under the Receiver Operating Characteristic curve (AUROC; Fawcett, 2006) and the Hosmer-Lemeshow goodness of fit (GOF) test [26] from '*ResourceSelection*' [27]. We checked the model misclassification error and concordance using functions in the package '*InformationValue*' [28].

Finally, we fitted $M_{final}$ to a resampled dataset to check for class bias, which can arise when data contain different proportions of responses (e.g. more ones than zeroes). The full dataset had differing proportions of each response (e.g. 1s and 0s), so we drew equal proportions of each response (to a total of 70% of the original sample size of the least common response) to use as a training dataset for the restricted model ($M_R$) based on the predictors in $M_{final}$. The remaining data were reserved for validation of $M_R$.

### SI-1.10 Post-hoc power analysis

For cases in which no logistic regression outperformed the null model for a given response variable, we performed simulation-based *post hoc* power analysis to establish the power associated with the test. First, for a given model *M*, we resampled (with replacement) from our observed data, then fit a logistic regression with these data and the model parameters specified by model *M.* We then conducted a likelihood ratio test to compare this model with a null model for the same data. This procedure was repeated 10,000 times, and we calculated *post hoc* power as the proportion of these tests that were significant at $\alpha = 0.05$. Whether a particular iteration is significant can be understood as the outcome of a Bernoulli trial with probability *p*, where *p* is the power. The proportion found to be significant over 10,000 iterations allows us to approximate the true *p*.

### SI-1.11 Microclimate predictors for analyses at site G1

We set out to investigate the contribution of microclimate to host plant phenology and condition within site G1. To do so, we modelled the quadrat-level condition and phenophase indices (Main Text) as a function of quadrat-specific microclimatic temperature and moisture availability. Individual quadrat locations within the site will have experienced the same broader-scale, site-level patterns of temperature and precipitation but may differ in the microclimate that they experience.

To estimate the microclimatic variability within site G1, we deployed thermochron iButtons (DS1921G-F5; Maxim Integrated, California, USA) to monitor the ground-level temperature at each of the 31 quadrat locations, at half-hourly intervals between 7$^{th}$ September to 8$^{th}$ October 2017. We used the maximum, mean and minimum temperatures as a relative measure of thermal microclimates across quadrats within the site. We used an HH2 moisture meter with ML3 ThetaProbe (Delta-t Devices, Cambridge, UK) to measure the soil moisture in each quadrat in June 2017. The average of three readings taken at random points within each quadrat was taken to represent quadrat soil moisture. All readings were taken within a 90 minute period of dry weather to minimise the effects of temporal variation in soil moisture.

Plant, temperature and moisture measurements were taken on different dates; we therefore consider the temperature and moisture measurements as indicators of microclimatic differences between quadrats which may be associated with host plant characteristics, rather than linking specific temperature or moisture regimes to cues or drivers of plant phenology or condition.

### SI-1.12 Supplementary Results

*SI-1.12.1 Inter-specific comparisons of host condition*

There was a substantial decline in quadrat-level condition of the annual host *G. dissectum* over summer 2016 that was not observed in the perennial *Helianthemum* or to the same extent in *Erodium* (Figure 2a; Table S4). However, by early November 2016, senesced *Geranium* had mostly been replaced by recently-germinated, better condition recruits, and the quadrat-level condition was at least as high as that of *Helianthemum* (Figure 2; Table S4).

Table S4. Summary of Kruskal-Wallis (KW) tests of differences in condition among host plants during each plant survey (July 2016–April 2017). Also presented, for all surveys with significant KW tests, are Dunn's tests (DT) of multiple comparisons between *Erodium cicutarium* (E), *Geranium dissectum* (G) and *Helianthemum nummularium* (H) species pairs. All *p* values are reported as adjusted ($p_{adj}$) with the Bonferroni correction, with a significance threshold set at 0.05; *** denotes $p_{adj}$ < 0.001.

| | | **Survey period** | | | | | | | | | |
|---|---|---|---|---|---|---|---|---|---|---|---|
| **Test** | **Test stat.** | **Jul. 2016** | **Aug. 2016** | **Sep. 2016** | **Oct. 2016** | **Nov. 2016** | **Dec. 2016** | **Jan. 2017** | **Feb. 2017** | **Mar. 2017** | **Apr. 2017** |
| **KW test** | **$X^2$ (df)** | 37.48 (2) | 111.2 (2) | 149.1 (2) | 69.87 (2) | 0.866 (2) | 6.828 (1) | 5.428 (1) | 3.787 (2) | 3.934 (2) | 0.571 (2) |
| | ***p*** | *** | *** | *** | *** | 0.649 | 0.009 | 0.020 | 0.151 | 0.140 | 0.742 |
| **DT: E/G** | ***Z*** | 0.834 | 6.051 | 7.555 | 5.517 | – | – | – | – | – | – |
| | **$p_{adj}$** | 0.202 | *** | *** | *** | – | – | – | – | – | – |
| **DT: E/H** | ***Z*** | -4.123 | -3.425 | -2.397 | -0.039 | – | – | – | – | – | – |
| | **$p_{adj}$** | *** | *** | 0.025 | 0.969 | – | – | – | – | – | – |
| **DT: G/H** | ***Z*** | -5.598 | -10.360 | -12.150 | -6.941 | – | 6.828 | 5.428 | – | – | – |
| | **$p_{adj}$** | *** | *** | *** | *** | – | 0.004 | 0.010 | – | – | – |

*SI-1.12.2 Inter-annual comparisons of Geraniaceae host characteristics*

There was temporal variation in host plant cover during each summer period (Figure S2), and higher cover of both Geraniaceae hosts in 2016 than 2017, except in October; Table S5). This suggests that, until October, there was a higher overall quantity of host available to brown argus in 2016 than 2017. Typically, this was because many plants in summer 2017 were small but good condition (and often newly germinated plants), whereas in 2016 the quadrats were dominated by fewer larger plants of more advanced phenology and poorer condition (JES *pers. obs.*).

The condition of the youngest *Geranium* in each quadrat was significantly higher in all 2017 survey periods compared to 2016 (Figure 2b; Table S5). *Erodium* showed similar patterns, though condition was approximately equivalent between years for the August and September surveys (Figure S3; Table S5).

*Geranium* phenophases differed significantly between years (Table S5): in 2016, the youngest plant in each *Geranium* quadrat was typically an older plant in the seed set stage (Figure 2c, main text) and new plants in the leaf stage did not dominate until October; however, this younger form was dominant throughout summer 2017 (Figure 2c, main text). *Erodium* quadrats were similarly dominated by young, leaf-stage plants in 2017: significantly more so than in 2016 during July and August (Figure S3; Table S5).

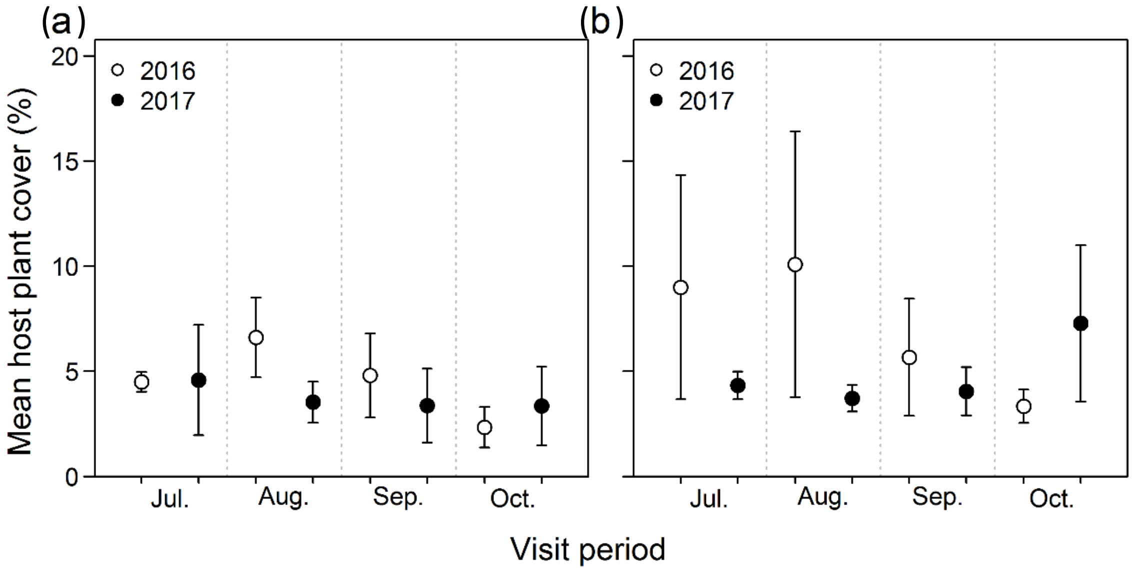

Figure S2. Mean (± SD) host plant ground cover (%) of (a) *Geranium dissectum* and (b) *Erodium cicutarium* across all sites visited during the summer (late July–early October) of both 2016 (open circles) and 2017 (filled circles).

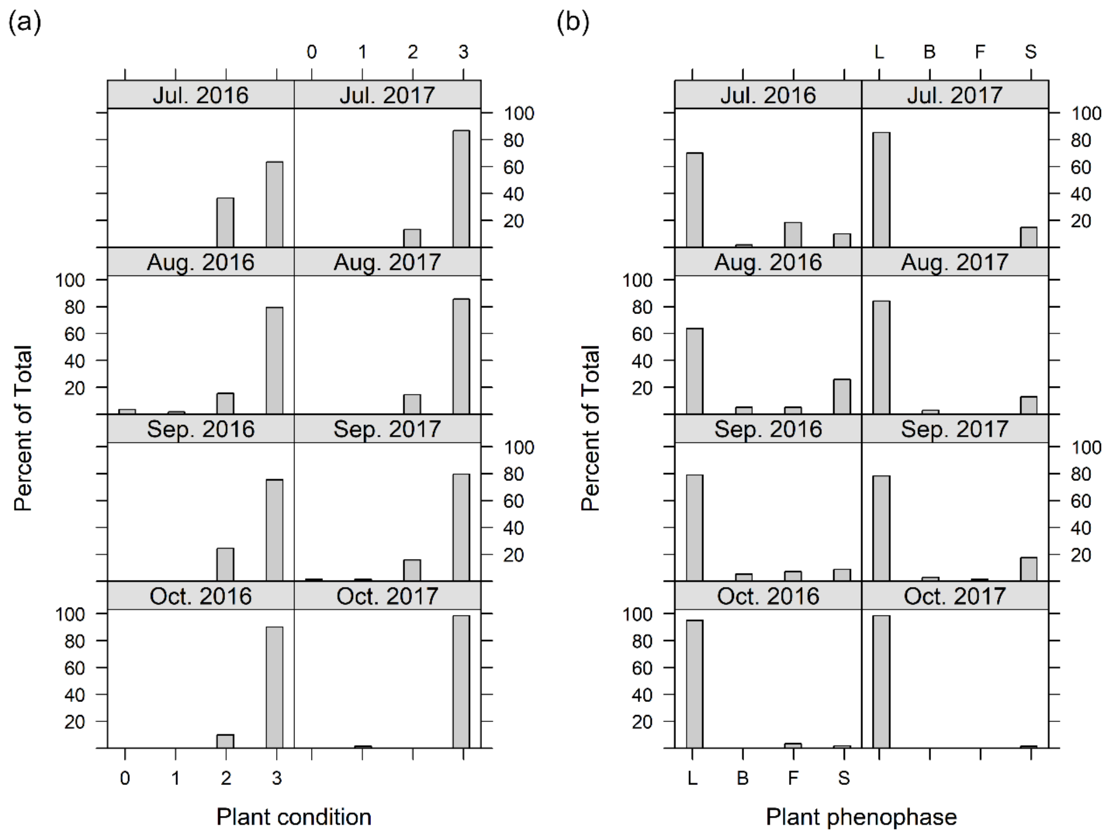

Figure S3. Condition (0–3; a) and phenophase (b) of the youngest *Erodium cicutarium* plant in each quadrat across all sites visited during late July–early October in both 2016 and 2017. Phenophase L: leaf; B: in bud; F: in flower; S: set seed.

Table S5. Summary of inter-annual comparisons (2016 versus 2017) of percentage cover, condition and phenophase for *Geranium dissectum* and *Erodium cicutarium* in all summer survey periods (July–October). Host cover and condition were compared using Mann-Whitney U (MWU) tests, host phenophases with chi-squared ($X^2$) tests of independence.

| Test | Test stat. | Survey period | | | |
|---|---|---|---|---|---|
| | | **July** | **August** | **September** | **October** |
| ***MWU: G. dissectum* percentage cover** | **W** | 8663.5 | 11796.5 | 8132.0 | 5667.0 |
| | ***p*** | < 0.001 | < 0.001 | < 0.001 | 0.652 |
| ***MWU: E. cicutarium* percentage cover** | **W** | 3945.5 | 4185.5 | 3436.0 | 2412.5 |
| | ***p*** | < 0.001 | < 0.001 | < 0.001 | 0.473 |
| ***MWU:* Quadrat-level *G. dissectum* condition** | **W** | 9147.0 | 4195.0 | 1655.5 | 2496.0 |
| | ***p*** | < 0.001 | < 0.001 | < 0.001 | < 0.001 |
| ***MWU:* Quadrat-level *E. cicutarium* condition** | **W** | 1540.5 | 1439.5 | 1556.0 | 1665.0 |
| | ***p*** | 0.003 | < 0.001 | 0.024 | 0.093 |
| ***MWU:* Condition of youngest *G. dissectum*** | **W** | 3697.0 | 1805.0 | 987.0 | 3538.0 |
| | ***p*** | <0.001 | 0.001 | 0.001 | 0.001 |
| ***MWU:* Condition of youngest *E. cicutarium*** | **W** | 1562.0 | 1901.5 | 1846.5 | 1734.0 |
| | ***p*** | 0.002 | 0.330 | 0.528 | 0.049 |
| **$X^2$ test: Quadrat-level *G. dissectum* phenophase** | ***n*** | 229 | 242 | 218 | 198 |
| | **$X^2$** | 10.714 | 50.158 | 76.389 | 53.536 |
| | ***p*** | 0.005 | 0.001 | 0.001 | 0.001 |
| **$X^2$ test: Quadrat-level *E. cicutarium* phenophase** | ***n*** | 127 | 129 | 126 | 123 |
| | **$X^2$** | 26.850 | 22.607 | 19.730 | 12.523 |
| | ***p*** | 0.001 | 0.001 | 0.001 | 0.006 |
| **$X^2$ test: Phenophase of youngest *G. dissectum*** | ***n*** | 229 | 232 | 216 | 199 |
| | **$X^2$** | 104.997 | 151.231 | 156.787 | 29.424 |
| | ***p*** | 0.001 | 0.001 | 0.001 | 0.001 |
| **$X^2$ test: Phenophase of youngest *E. cicutarium*** | ***n*** | 128 | 127 | 126 | 123 |
| | **$X^2$** | 15.119 | 8.453 | 4.599 | 2.138 |
| | ***p*** | 0.002 | 0.038 | 0.204 | 0.343 |

*SI-1.12.3 Quadrat-level variation in host characteristics*

The Geraniaceae host plants showed intra- and inter-annual variability in condition and phenology over consecutive summers. Specifically, quadrat-level classifications suggest that *G. dissectum* shows marked intra-annual changes and inter-annual differences in condition and phenology: Mann-Whitney U tests confirmed that quadrat-level condition was higher in 2017 than 2016 in all summer surveys except July (Figure S4; Table S5), and Chi-squared tests confirmed that the availability of host phenophases differed significantly between 2016 and 2017 (Table S5), suggesting higher availability of young pre-reproductive plants in 2017 (Figure S5).

*E. cicutarium* also shows intra- and inter-annual variation in both condition and phenology: quadrat-level condition was marginally higher in 2017 for all surveys except October (Figure S4; Table S5), and 2017 was characterised by higher availability of younger, pre-reproductive plants than in 2016, which had higher proportions of quadrats categorised as being in bud, flower or seed-setting phenophases (Figure S5; Table S5).

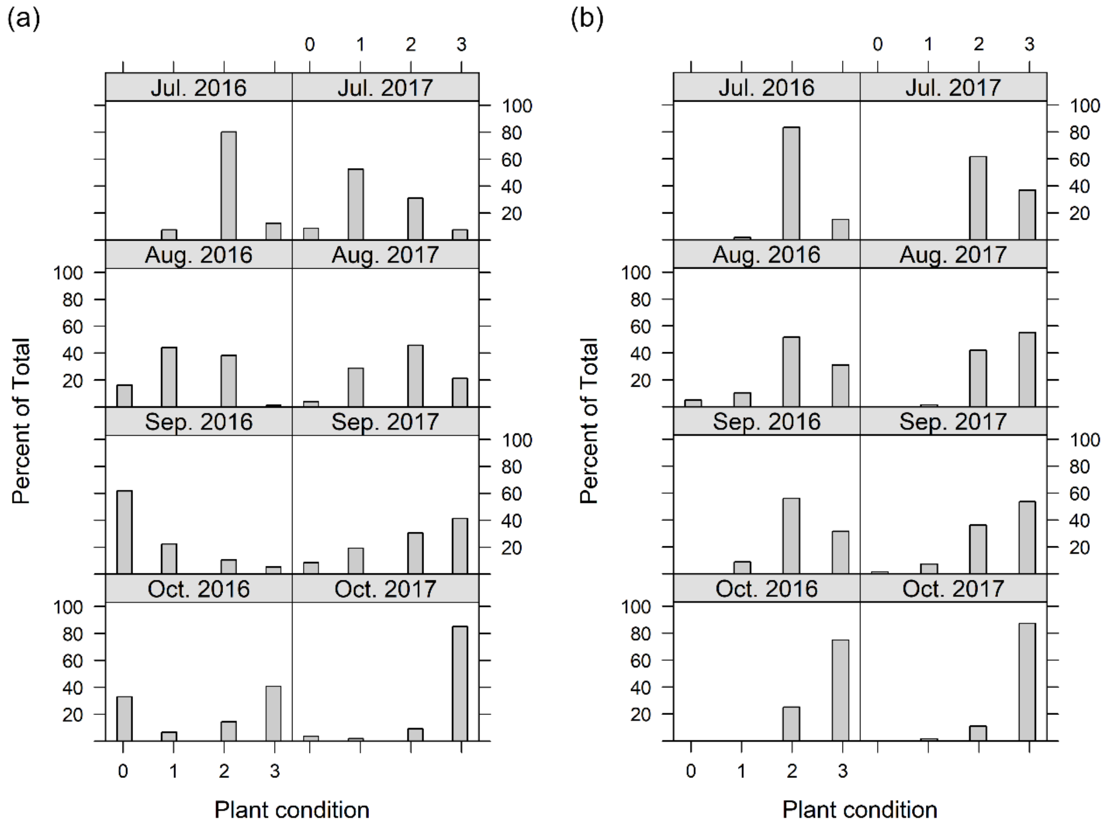

Figure S4. Quadrat-level condition (0–3) of (a) *Geranium dissectum* and (b) *Erodium cicutarium* was typically higher across all sites visited during summer (late July–early October) 2017 than summer 2016.

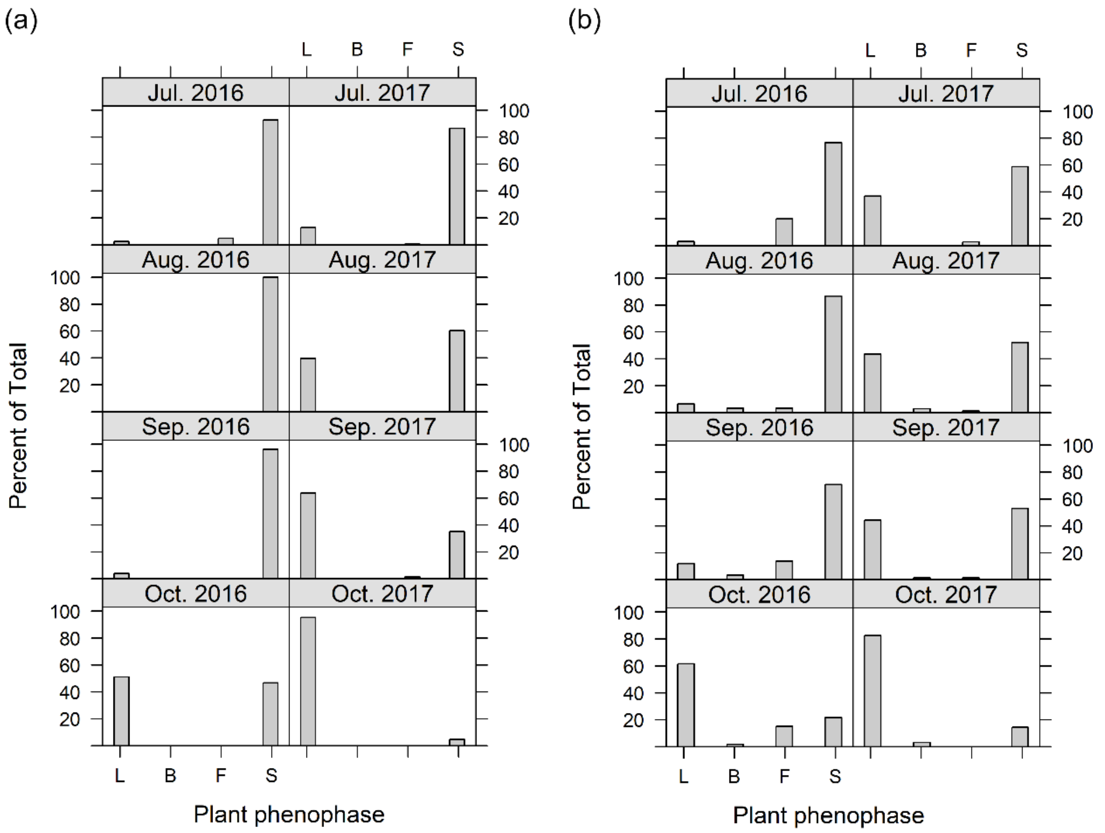

Figure S5. Quadrat-level categorisations of the phenophases of (a) *Geranium dissectum* and (b) *Erodium cicutarium* across all visited sites during the summer (late July–early October) of 2016 and 2017. Phenophase L: leaf; B: in bud; F: in flower; S: set seed. There was higher availability of younger, pre-reproductive plants in summer 2017 than summer 2016.

*SI-1.12.4 Plant-herbivore (a)synchrony*

We assessed potential for asynchrony between the butterfly and its hosts (question (b)) by overlaying plots of site-specific host condition indices with curves representing brown argus phenology (Figure 3), and performing beta regressions on derived asynchrony estimates. The beta regressions of AUC synchrony estimates demonstrate that potential asynchrony was highest for brown argus on *Geranium* in 2016 (low AUC overlap: AUC range 0.51–0.73), but very low for other hosts and for *Geranium* in 2017 (high AUC overlap: AUC range 0.92–1.00) (Figure 3, Table S6). Potential asynchrony was more pronounced for larvae than adults, especially on *Geranium* in 2016 (Table S6; Figures 3 and S6). There was no detectable effect of site latitude on adult or larval AUC overlap: it was not possible to fit models with site latitude as a predictor, but a basic follow-up analysis also showed no correlation between the AUC overlap estimates and site latitude for either adult or larval brown argus (Spearman's $\rho < 0.041$, $p > 0.88$). All diagnostic tests for Bayesian beta regressions were acceptable, producing a models with conditional $R^2$ of 0.851 (89% CI [0.685, 0.897]) for adults and 0.882 (89% CI [0.740, 0.922]) for larvae, good Pareto k estimates and convergence for all parameters. The beta regressions (Table 1) showed a clear host-year interaction effect such that the plant-host synchrony was lowest on Geranium in 2016 than in 2017 or on other hosts (Table 1).

Table S6. Adult and larval area under the curve (AUC) synchrony estimates for each site-year combination, each calculated as the full area of the phenology curve minus the area which lies above the corresponding host condition line

| | | 2016 | | 2017 | |
|---|---|---|---|---|---|
| **Host** | **Site** | **Adult AUC** | **Larval AUC** | **Adult AUC** | **Larval AUC** |
| *Geranium* | G1 | 0.728 | 0.644 | 0.921 | 0.922 |
| *Geranium* | G2 | 0.767 | 0.707 | 1.00 | 1.00 |
| *Geranium* | G3 | 0.599 | 0.509 | 1.00 | 1.00 |
| *Erodium* | E1 | 0.982 | 0.983 | 0.982 | 0.976 |
| *Erodium* | E2 | 0.972 | 0.977 | 0.994 | 0.993 |
| *Helianthemum* | H1 | 0.954 | 0.963 | – | – |
| *Helianthemum* | H2 | 0.944 | 0.945 | – | – |
| *Helianthemum* | H3 | 0.927 | 0.929 | – | – |
| *Helianthemum* | H4 | 0.986 | 0.978 | – | – |
| *Helianthemum* | H5 | 0.987 | 0.979 | – | – |

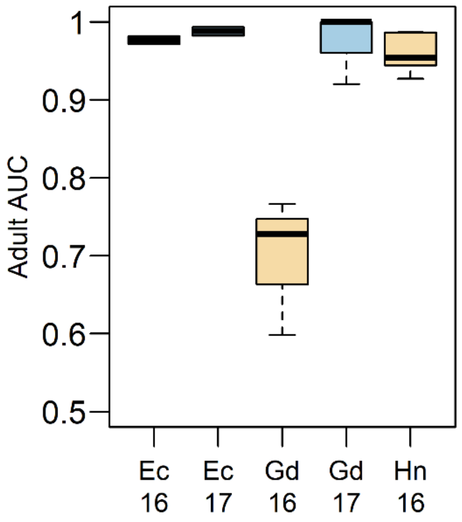

Figure S6. Summary of adult AUC synchrony metrics for each host-year combination, summarised from site-specific metrics each calculated as the full area of the phenology curve minus that which lies above the corresponding host condition line. The equivalent plot for larvae is shown in Figure 3.

#### *SI-1.12.5 Drivers of late summer recruitment and condition of Erodium cicutarium and Geranium dissectum*

Model selection based on logistic regressions (binomial GLMs) suggested that no models out-performed a null model for the probability of new recruitment of *E. cicutarium* within a quadrat, though summer rainfall may be influential (Table S7). However, simulation-based *post hoc* power analysis indicate that we had low power (0.52) to detect a significant effect based on the available data and the model parameters reported for model $M_a$ regarding the probability of new recruitment (Table S7). Only 13% of simulations of a model with the same structure as $M_a$ had an AIC better than the AIC of the null model.

The best AIC model for condition of the youngest *E. cicutarium* plant within each quadrat was the null model, and there were no competing models (data not shown). However, simulation-based *post hoc* power analysis indicated that we had very low power (0.27) to detect an effect, and only 4% of simulations resulted in a model with an AIC better than the null model. The power analysis of the binomial GLM for the condition of the youngest *E. cicutarium* plant was based on the model parameters associated with the (rejected) model that had the closest AIC value to that of the null model. July to September rainfall was the only variable in this rejected model. Taken together, these results for *E. cicutarium* may reflect the relatively consistently good condition of *E. cicutarium* observed across years, sites and weather conditions (Figure S4) or a lack of sufficient variability in climatic data from these sites to detect a pattern.

Tables 1 (main text), S7 and S8 include parameter estimates for the 'Site' terms (S), which were code as 'sum contrasts', as outlined in SI-1.9 above.

Table S7. Drivers of the probability of new recruitment of *Erodium cicutarium* during September surveys (2016–2017), summarising AIC analyses and parameter estimates (with standard errors) of candidate models ($M_a$, $M_b$,…$M_z$) with ΔAIC ≤ 6. $M_{best}$ denotes the best AIC model and $M_{final}$ denotes the selected, most parsimonious model: in this case the null model (elsewhere denoted $M_N$). All selected models contain a subset of fixed effects from: daily mean rainfall (R) in either July (J) or the late summer period (L) between August 1st–sampling date in September. Other terms were tested but were not detected in the final candidate set of models with ΔAIC ≤ 6. $\beta_0$ is the intercept, *k* is the number of parameters and *LL* is the log-likelihood of the model.

| | **Model parameters** | | | | | |
|---|---|---|---|---|---|---|
| **Model** | $\beta_0$ | $R_J$ | $R_L$ | ***k*** | ***LL*** | **ΔAIC** |
| $M_{best}$ | -0.065 (0.630) | 1.152 (0.231) | -3.706 (1.304) | 3 | -66.77 | 0.00 |
| $M_a$ | 0.200 (0.633) | – | -2.091 (1.333) | 2 | -67.92 | 0.30 |
| $M_{final}$ | 1.163 (0.209) | – | – | 1 | -69.16 | 0.78 |

Nested models are shown only if the inclusion of additional parameters improves the AIC value for that model relative to the simpler version.

The probability of the youngest *Geranium* plants being in good condition increased with summer rainfall ($MC_{final}$; Figure 4a, Table 2a, Table S8). For $MC_{final}$, a Hosmer-Lemeshow goodness-of-fit test provides no reason to suspect a lack of fit ($X^2$ = 1.246, df = 8, $p$ = 0.996), and the area under the Receiver Operating Characteristic (ROC) curve (AUROC = 0.939) suggests the model has high predictive ability. This model also has high concordance (90.24%), and low misclassification rate (11%). The restricted model ($MC_R$; constructed to check for class bias) returned similar parameter estimates to the full model (Table 2, Main Text). The AUROC (0.943) and concordance (0.907) also indicate excellent predictive ability of $M_R$ against test data, for which there was a very low (8%) misclassification rate. Forcing models of *Geranium* condition to include an effect of year resulted in higher AIC values and/or inflated the estimates and standard errors by several orders of magnitude, so these are not reported in the main text, and there are no effects of year in the final model set. However, models with an effect of year are shown in Table S8 for context. The candidate model for *Geranium* condition (*MC*) that include year effects but that do not suffer from inflated parameter estimates and standard errors (Table S8, model $MC_f$) instead has a high ΔAIC value, suggesting a lack of parsimony relative to the final model [23,24].

The probability of new *Geranium* recruitment was higher in 2017 ($MR_{best}$, Table S9). This model was not chosen as the final model ($MR_{final}$) as it was considered more informative to also account for the known climatic differences between the years (which contribute to the between-year differences in recruitment). Therefore, there is evidence to suggest that the probability of new *Geranium* recruitment was lower following higher mean daily temperatures during August–September ($MR_{final}$; Figure 4b, Table 2b, Table S9), reflecting the lower August-September temperatures in 2017 than 2016. A Hosmer-Lemeshow goodness-of-fit test provides no reason to suspect a lack of fit for $MR_{final}$ ($X^2$ = 6.111, df = 8, $p$ = 0.635), and the area under the Receiver Operating Characteristic (ROC) curve (AUROC = 0.961) suggests the model has high predictive ability. This model also has high concordance (93.33%), and low misclassification rate (7%). The restricted model ($MR_R$) constructed to check for class bias returned similar parameter estimates to the full model (Table 2). The AUROC (0.971) and concordance (0.946) also indicate excellent predictive ability of $M_R$ against test data, for which there was a very low (4%) misclassification rate.

Forcing $MR_{final}$ (Tables 1 and S8) to include an effect of year (as in model $M_a$, Table S9) resulted in inflated parameter estimates and standard errors, likely due to high collinearity between the variables. Therefore, these are not reported in the main text, but are reported in Table S9 for context.

Table S8. Drivers of the condition of the youngest *Geranium dissectum* in each quadrat during September surveys (2016–2017), summarising AIC analyses and parameter estimates (with standard errors) for all candidate logistic regression models ($M_a$, $M_b$,…$M_z$) with ΔAIC ≤ 6. $M_{best}$ denotes the best AIC model and $M_{final}$ denotes the selected, most parsimonious model. Also presented for comparison are the null model ($M_N$) and a restricted model ($M_R$) used to check for class bias in $M_{final}$. All selected models contain a subset of fixed effects from: site ($S_1$ and $S_2$), daily mean rainfall (R) or aridity (A) in either July ($_J$), the summer period ($_S$) between July 1st–sampling date in September or the late summer period ($_L$) between August 1st–sampling date in September, mean vegetation height (*V*), and year (*Y*). Other terms were tested but were not detected in the final candidate set of models with ΔAIC ≤ 6. $\beta_0$ is the intercept, which accounts for the mean of all site effects (each represented by $S_1$ and $S_2$ as zero-sum contrasts) in all but the null model ($M_N$), *k* is the number of parameters and *LL* is the log-likelihood of the model.

| | Model parameters | | | | | | | | | | | | |
|---|---|---|---|---|---|---|---|---|---|---|---|---|---|
| **Model** | **$\beta_0$** | $S_1$ | $S_2$ | $R_S$ | $R_L$ | $A_J$ | $A_S$ | $A_L$ | $V$ | $Y$ | ***k*** | ***LL*** | **ΔAIC** |
| **$MC_{best}$** | 1.621<br>(0.393) | 1.464<br>(0.413) | -0.352<br>(0.384) | 3.627<br>(0.509) | – | – | – | – | -0.728<br>(0.290) | – | 5 | -60.11 | 0.00 |
| **$MC_{final}$** | 2.053<br>(0.535) | 5.749<br>(0.985) | -1.590<br>(0.551) | – | 5.871<br>(0.891) | – | – | – | – | – | 4 | -61.17 | 0.13 |
| **$MC_a$†** | 11.687<br>(807.169) | 28.224<br>(1863.9) | -9.204<br>(647.974) | – | 27.693<br>(1777.7) | – | – | – | – | 5.423<br>(424.726) | 5 | -60.20 | 0.20 |
| **$MC_b$** | 1.566<br>(0.377) | 1.616<br>(0.397) | -0.497<br>(0.388) | – | – | – | 3.558<br>(0.496) | – | -0.700<br>(0.285) | – | 5 | -61.43 | 2.64 |
| **$MC_c$** | 1.562<br>(0.376) | 1.015<br>(0.386) | -0.176<br>(0.374) | – | – | 3.491<br>(0.487) | – | – | -0.700<br>(0.285) | – | 5 | -61.45 | 2.68 |
| **$MC_d$** | 1.471<br>(0.364) | 4.069<br>(0.578) | -1.781<br>(0.479) | – | – | – | – | 4.096<br>(0.568) | -0.695<br>(0.284) | – | 5 | -61.72 | 3.22 |
| **$MC_e$** | 1.602<br>(0.398) | 1.445<br>(0.424) | 0.098<br>(0.347) | 3.728<br>(0.508) | – | – | – | – | – | – | 4 | -63.57 | 4.92 |
| **$MC_f$†** | 0.309<br>(0.225) | 0.450<br>(0.330) | -0.688<br>(0.320) | – | – | – | – | – | – | -2.135<br>(0.242) | 4 | -72.39 | 22.57 |
| **$MC_N$** | 0.545<br>(0.141) | – | – | – | – | – | – | – | – | – | 1 | -143.30 | 168.38 |
| **$MC_R$** | 1.565<br>(0.622) | 5.071<br>(1.116) | -1.305<br>(0.630) | – | 5.611<br>(1.128) | – | – | – | – | – | 4 | * | * |

* We do not report the log-likelihood or ΔAIC for $MC_R$ because this model was tested on a subset of data and so is not comparable in these terms.
Nested models are shown only if the inclusion of additional parameters improves the AIC value for that model relative to the simpler version.
†Models forced to include year effects, which show inflated parameter estimates and standard errors, or large AIC values.

Table S9. Drivers of the probability of new recruitment of *Geranium dissectum* during September surveys (2016–2017), summarising AIC analyses and parameter estimates (with standard errors) for all candidate logistic regression models ($M_a$, $M_b$,…$M_z$) with ΔAIC ≤ 6. $M_{best}$ denotes the best AIC model and $M_{final}$ denotes the selected, most parsimonious model. Also presented for comparison are the null model ($M_N$) and a restricted model ($M_R$) used to check for class bias in $M_{final}$. All selected models contain a subset of fixed effects from: site ($S_1$ and $S_2$), mean daily temperature in the late summer period ($T_L$) between August 1st–sampling date in September, mean vegetation height ($V$), and year ($Y$). Other terms were tested but were not detected in the final candidate set of models with ΔAIC ≤ 6. $\beta_0$ is the intercept accounting for the mean of all site effects (each represented by $S_1$ and $S_2$ as zero-sum contrasts), $k$ is the number of estimated parameters and $LL$ is the log-likelihood of the model.

| | Model parameters | | | | | | | | |
|---|---|---|---|---|---|---|---|---|---|
| **Model** | $\beta_0$ | $S_1$ | $S_2$ | $T_L$ | $V$ | $Y$ | ***k*** | ***LL*** | **ΔAIC** |
| **M**$R_a$† | -467.6 (36163.3) | -335.3 (27017.8) | 178.0 (14325.9) | 364.1 (27824.4) | – | 717.8 (54570.0) | 5 | -38.84 | -17.04 |
| **M**$R_{best}$† | -3.984 (0.753) | 1.074 (0.577) | -1.627 (0.520) | – | – | 6.898 (0.989) | 4 | -46.85 | 0.00 |
| **M**$R_b$ | 0.289 (0.369) | 4.482 (0.817) | -3.968 (0.784) | -3.862 (0.570) | -0.626 (0.347) | – | 4 | -47.36 | 3.01 |
| **M**$R_{final}$ | 0.310 (0.354) | 4.366 (0.794) | -3.558 (0.718) | -3.872 (0.555) | – | – | 4 | -49.07 | 4.44 |
| **M**$R_N$ | 0.352 (0.138) | – | – | – | – | – | 1 | -147.78 | 192.84 |
| **M**$R_R$ | -0.044 (0.495) | 4.770 (1.126) | -4.120 (1.017) | -4.096 (0.792) | – | – | 4 | * | * |

* We do not report the log-likelihood or ΔAIC for $M_R$ because this model was tested on a subset of data and so is not comparable in these terms.
Nested models are shown only if the inclusion of additional parameters improves the AIC value for that model relative to the simpler version.
†Models with year effects. M$R_a$ shows inflated, unreliable parameter estimates and standard errors, likely due to collinearity between year and weather variables. This model was therefore not considered to represent the best AIC model (M$R_{best}$).

*SI-1.12.6 Testing for effects of microclimate on Geranium condition and recruitment*

At site G1 in September 2017, areas of moister soil (where plants are less likely to dry out) were associated with *Geranium* plants in better condition and earlier phenophases (Figure 4c,d and Table 3, main text). Candidate models suggested such plants were also more prevalent in areas with warmer, moister microclimates (Tables S9 and S10), though quadratic effects suggest a threshold beyond which increasing temperature may be detrimental to host condition even in moist microclimates. Candidate models with only temperature effects did not outperform the null model (Table S10 and S11).

Table S10. Summary of AIC analyses and parameter estimates (with standard errors) for variables associated with within-site quadrat-specific condition indices of *Geranium dissectum* at site G1 (Bubbenhall Meadow) in 2017. All models ($M_a$, $M_b$,…$M_z$) with ΔAIC ≤ 6 are presented; $M_{best}$ denotes the best AIC model, and the selected 'final' model is denoted $M_{final}$. Also presented for comparison is the null model ($M_N$). All selected models contain a subset of fixed effects from: soil moisture (M), mean (T) and minimum temperature (t). Other terms and interactions were tested but were not detected in the final candidate set of models with ΔAIC ≤ 6. $\beta_0$ is the intercept, *k* is the number of parameters and *LL* is the log-likelihood of the model.

| Model | Model parameters | | | | *k* | *LL* | ΔAIC |
|---|---|---|---|---|---|---|---|
| | $\beta_0$ | M | T | t | | | |
| **M*CI*$_{best}$** | 0.586 (0.453) | 0.602 (0.511) | 1.172 (0.690) | -0.816 (0.600) | 4 | -12.65 | 0.00 |
| **M*CI*$_{final}$** | 0.496 (0.413) | 0.810 (0.448) | – | – | 2 | -15.59 | 1.88 |
| **M*CI*$_a$** | 0.550 (0.435) | – | 1.387 (0.658) | -0.756 (0.577) | 3 | -15.03 | 2.75 |
| **M*CI*$_b$** | 0.497 (0.413) | – | 0.867 (0.489) | – | 2 | -17.27 | 5.23 |
| **M*CI*$_N$** | 0.428 (0.380) | – | – | – | 1 | -19.69 | 8.07 |

Nested models are shown only if the inclusion of additional parameters improves the AIC value for that model relative to the simpler version.

Table S11. Summary of AIC analyses and parameter estimates (with standard errors) for variables associated with the within-site quadrat-level phenophase indices of *Geranium dissectum* at site G1 in 2017. All models ($M_a$, $M_b$,…$M_z$) with ΔAIC ≤ 6 are presented; $M_{best}$ denotes the best AIC model, and the selected 'final' model is denoted $M_{final}$. Also presented for comparison is the null model ($M_N$). All selected models contain a subset of fixed effects from: bare ground cover (G), soil moisture (M), mean temperature (T) and minimum temperature (t). Other terms and interactions were tested but were not detected in the final candidate set of models with ΔAIC ≤ 6. $\beta_0$ is the intercept, *k* is the number of parameters and *LL* is the log-likelihood of the model.

| Model | Model parameters | | | | | *k* | *LL* | ΔAIC |
|---|---|---|---|---|---|---|---|---|
| | $\beta_0$ | *G* | *M* | *T* | *t* | | | |
| **M*PI*$_{best}$** | 0.642 (0.579) | 1.732 (1.576) | 0.889 (0.446) | – | – | 3 | -13.06 | 0.00 |
| **M*PI*$_a$** | 0.455 (0.456) | – | 0.641 (0.514) | 1.298 (0.712) | -0.823 (0.613) | 4 | -12.52 | 0.92 |
| **M*PI*$_{final}$** | 0.366 (0.412) | – | 0.889 (0.453) | – | – | 2 | -14.84 | 1.57 |
| **M*PI*$_b$** | 0.423 (0.437) | – | – | 1.524 (0.680) | -0.747 (0.585) | 3 | -15.45 | 4.78 |
| **M*PI*$_N$** | 0.307 (0.376) | – | – | – | – | 1 | -19.98 | 9.85 |

Nested models are shown only if the inclusion of additional parameters improves the AIC value for that model relative to the simpler version.

### References

1. Pollard E, Hall ML, Bibby TJ. 1986 Monitoring the abundance of butterflies 1976-1985.

2. UKBMS. 2017 UK Butterfly Monitoring Scheme (UKBMS) occurrence dataset https://doi.org/10.15468/gmqvmk accessible via GBIF.org.

3. Heath JH, Pollard E, Thomas JA. 1984 *Atlas of Butterflies in Britain and Ireland*. Harmondsworh, United Kingdom: Viking. See https://www.abebooks.co.uk/9780670800063/Atlas-Butterflies-Britain-Ireland-Heath-0670800066/plp.

4. Lakhani KH, Davis BNK. 1982 Multiple regression models of the distribution of *Helianthemum chamaecistus* in relation to aspect and slope at Barnack, England. *Journal of Applied Ecology* **19**, 621–629.

5. Bridle JR, Buckley J, Bodsworth EJ, Thomas CD. 2014 Evolution on the move: specialization on widespread resources associated with rapid range expansion in response to climate change. *Proceedings of the Royal Society B: Biological Sciences* **281**, 20131800. (doi:10.1098/rspb.2013.1800)

6. Asher J, Warren MS, Fox R, Harding P, Jeffcoate G, Jeffcoate S. 2001 *The Millennium Atlas of Butterflies In Britain and Ireland*. Oxford, United Kingdom: Oxford University Press.

7. Suggitt AJ, Gillingham PK, Hill JK, Huntley B, Kunin WE, Roy DB, Thomas CD. 2011 Habitat microclimates drive fine-scale variation in extreme temperatures. *Oikos* **120**, 1–8. (doi:10.1111/j.1600-0706.2010.18270.x)

8. Bramer I *et al.* 2018 Advances in monitoring and modelling climate at ecologically relevant scales. *Next Generation Biomonitoring, part 1* **58**, 101–161. (doi:http://dx.doi.org/10.1016/bs.aecr.2017.12.005)

9. Bourn NAD, Thomas JA. 1993 The ecology and conservation of the brown argus butterfly *Aricia agestis* in Britain. *Biological Conservation* **63**, 67–74. (doi:10.1016/0006-3207(93)90075-C)

10. Stewart JE, Maclean IMD, Edney EJ, Bridle JR, Wilson RJ. 2021 Microclimate and resource quality determine resource use in a range-expanding herbivore. *Biology Letters* **17**, 20210175. (doi:10.1098/rsbl.2021.0175)

11. Garroutte EL, Hansen AJ, Lawrence RL. 2016 Using NDVI and EVI to map spatiotemporal variation in the biomass and quality of forage for migratory elk in the Greater Yellowstone ecosystem. *Remote Sensing* **8**, 404. (doi:10.3390/rs8050404)

12. Wang Q, Tenhunen J, Dinh NQ, Reichstein M, Vesala T, Keronen P. 2004 Similarities in ground- and satellite-based NDVI time series and their relationship to physiological activity of a Scots pine forest in Finland. *Remote Sensing of Environment* **93**, 225–237. (doi:10.1016/j.rse.2004.07.006)

13. Dennis EB, Morgan BJT, Freeman SN, Roy DB, Brereton T. 2016 Dynamic models for longitudinal butterfly data. *JABES* **21**, 1–21. (doi:10.1007/s13253-015-0216-3)

14. Stewart JE. 2020 Linking Phenology to Population Dynamics and Distribution Change in a Changing Climate. PhD Thesis, University of Exeter.

15. Pollard E, Yates TJ. 1993 *Monitoring Butterflies for Ecology and Conservation: The British Butterfly Monitoring Scheme*. Springer Netherlands.

16. Met Office. 2017 UKCP09: Met Office gridded land surface climate observations - daily temperature and precipitation at 5km resolution. Updated to 2017.

17. Bürkner P-C. 2017 brms: An R Package for Bayesian Multilevel Models Using Stan. *Journal of Statistical Software* **80**, 1–28. (doi:10.18637/jss.v080.i01)

18. Hartig F. 2018 *DHARMa: Residual Diagnostics for Hierarchical (Multi-Level / Mixed) Regression Models. R package version 0.2.0.* See https://CRAN.R-project.org/package=DHARMa.

19. Gabry J, Mahr T. 2021 *bayesplot: Plotting for Bayesian Models. R package version 1.8.0.* See https://mc-stan.org/bayesplot/.

20. Lüdecke D, Ben-Shachar MS, Patil I, Waggoner P, Makowski D. 2021 performance: An R Package for Assessment, Comparison and Testing of Statistical Models. (doi:10.31234/osf.io/vtq8f)

21. Fox J, Weisberg. 2019 *An R Companion to Applied Regression*. Third Edition. Thousand Oaks, CA, U.S.A.: Sage.

22. R Core Team. 2018 *R: A language and environment for statistical computing.* Vienna, Austria: R Foundation for Statistical Computing. See https://www.R-project.org/.

23. Richards SA. 2008 Dealing with overdispersed count data in applied ecology. *Journal of Applied Ecology* **45**, 218–227. (doi:10.1111/j.1365-2664.2007.01377.x)

24. Richards SA. 2015 Likelihood and model selection. In *Ecological Statistics: Contemporary Theory and Application* (eds S Negrete-Yankelevich, VJ Sosa), pp. 58–80. Oxford, United Kingdom: Oxford University Press.

25. Mundry R, Nunn CL. 2009 Stepwise model fitting and statistical inference: turning noise into signal pollution. *The American Naturalist* **173**, 119–123. (doi:10.1086/593303)

26. Fagerland MW, Hosmer DW. 2016 Tests for goodness of fit in ordinal logistic regression models. *Journal of Statistical Computation and Simulation* **86**, 3398–3418. (doi:10.1080/00949655.2016.1156682)

27. Lele SR, Keim JL, Solymos P. 2017 *ResourceSelection: Resource Selection (Probability) Functions for Use-Availability Data. R Package version 0.3.2*. See https://cran.r-project.org/web/packages/ResourceSelection/index.html.

28. Prabhakaran S. 2016 *InformationValue: Performance Analysis and Companion Functions for Binary Classification Models. R Package version 1.2.3*. See https://cran.r-project.org/web/packages/InformationValue/index.html.
