## Supplementary material for "Climate-driven variation in biotic interactions provides a narrow and variable window of opportunity for an insect herbivore at its ecological margin": Summary of supplementary material

ESM 1: Site information and survey dates, Supplementary methods and Supplementary results, including additional figures and tables as outlined in the main text.

ESM 2: R code for all analyses and plotting

Provides all information including README information required for data management, analyses and plotting.

ESM 3: Geranium data

Contains data on the *Geranium dissectum* host plants surveyed throughout this study

Use of these data is highlighted in the attached R code, which uses relevant columns from this .csv file.

ESM 4: Erodium data

Contains data on the *Erodium cicutarium* host plants surveyed throughout this study

Use of these data is highlighted in the attached R code, which uses relevant columns from this .csv file.

ESM 5: Site G1 data

Contains data for the microclimate and host plant assessments and analyses from site G1 in 2017.

Use of these data is highlighted in the attached R code, which uses relevant columns from this .csv file.

ESM 6 & ESM 7: Summer quadrat data and 'All quadrat data' contain data on all host plant surveys.

Use of these data is highlighted in the attached R code, which uses relevant columns from this .csv file.
